# Music-Inspired Acoustic-Piezoelectric Stimulation Accelerates Extracellular Vesicle Production and Programs Therapeutic Function

**DOI:** 10.64898/2026.03.11.711216

**Authors:** James Johnston, Erin Boyce, Tiago Thomaz Migliati Zanon, Hyunsu Jeon, Courtney Khong, Yun Young Choi, Nosang V. Myung, Martin Nunez, Mae-Lin Pinkstaff, Yichun Wang

## Abstract

Macrophage small extracellular vesicles (sEVs) carry phenotype-linked cargo and bioactivity for immunomodulation and regeneration, but therapeutic translation is limited by low secretion and poor control of function. We introduce a music-activated piezoelectric nanofiber substrate (PES) that converted audible sound into programmable electrical stimulation to enhance sEV biogenesis while tuning macrophage polarization. Adjusting acoustic parameters increased sEV yield, while musically inspired “assemblies” biased macrophage phenotypes: dissonant, low-frequency stimuli promoted M1-like inflammation, whereas consonant, higher-frequency stimuli favored M2-like, regenerative states. These shifts produced distinct sEV cargo and bioactivities. We rationally designed customized music stimulus that maximized both vesicle production and M2 bias, yielding sEVs exhibited regeneration potentials. This work establishes a programmable acoustic–piezoelectric strategy to scale macrophage sEV production while tailoring their therapeutic potency.

## 1. Introduction

Small extracellular vesicles (sEVs) are nature-derived, lipid-based nanoparticles that that mediate intercellular communication and are emerging as a versatile therapeutic modality for cancer therapy, immunomodulation, and tissue regeneration. [1–5] In particular, sEVs secreted by tissue-resident macrophages are attractive because their molecular cargo and resulting bioactivity are tightly coupled to the activation state of the parent cells, enabling phenotype-dependent modulation of immune responses and regenerative processes. [6,7] Yet, realizing the full therapeutic potential of macrophage-derived sEVs will require strategies that enhance secretion and enable tunable control over their native functions. [8,9]

Piezoelectricity is a pervasive feature of biological tissues and extracellular matrices (ECM), providing cells with dynamic electrical cues that influence mechanotransduction and cell state. [10,11] Inspired by this feature, piezoelectric materials have been leveraged to stimulate cells and enhance sEV production by converting mechanical inputs into electrical stimulation. [12–15] The resulting electrical outputs can be programmed through acoustic parameters such as frequency, amplitude, timbre, and harmonic structure, well established in audible acoustic sensing systems. [16,17] Hence, ECM–mimetic piezoelectric platforms may provide a unified strategy to both enhance sEV biogenesis and reprogram key cellular pathways that govern the composition and functional potency of the resulting sEVs.

Meanwhile, acoustic stimuli are biologically relevant signals for innate immune cells. Macrophages exhibit measurable responses to audible sound exposures, including shifts in cytokine expression associated with polarization state. [18,19] Prior studies report that music dominated by consonant, higher-frequency components (e.g., classical and opera) is associated with increased anti-inflammatory cytokines, aligning with regeneration phenotypes. [20,21] Conversely, intense or dysregulated stimulation can promote proinflammatory signaling, desirable in select immunotherapy contexts. [19,22,23] Related approaches, such as vibroacoustic therapies (VAT), use bulk low frequency waves (20-150 Hz) to modulate immune system through combined mechanical and bioelectrical pathways at the cellular level. [24–27] However, how music-theory features beyond basic acoustic parameters translate through the piezoelectric elements of native tissues and the ECM into bioelectrical cues that program innate immune system remains poorly understood, limiting the rational design of acoustic-piezoelectric regimens for immunomodulation and regenerative medicine.

Here we introduce a music-activated piezoelectric nanofiber substrate (PES) to transduce audible acoustic waves to cellular electrical stimulation, enabling regulation of macrophage-derived sEV production and macrophage polarization (**Fig. 1**). We show that PES activation significantly accelerate sEV production and that musically inspired acoustic “assemblies” can bias macrophage polarization to tune sEV bioactivity. Finally, we engineered a customized regenerative musical stimulus that jointly optimizes sEV production and regeneration function, yielding high sEV concentration and enhanced osteoblast proliferation and gap closure. Together, this music-inspired acoustic–piezoelectric platform provides a programmable strategy to scale sEV production while simultaneously tailoring therapeutic properties through phenotype control.

**Figure 1.**
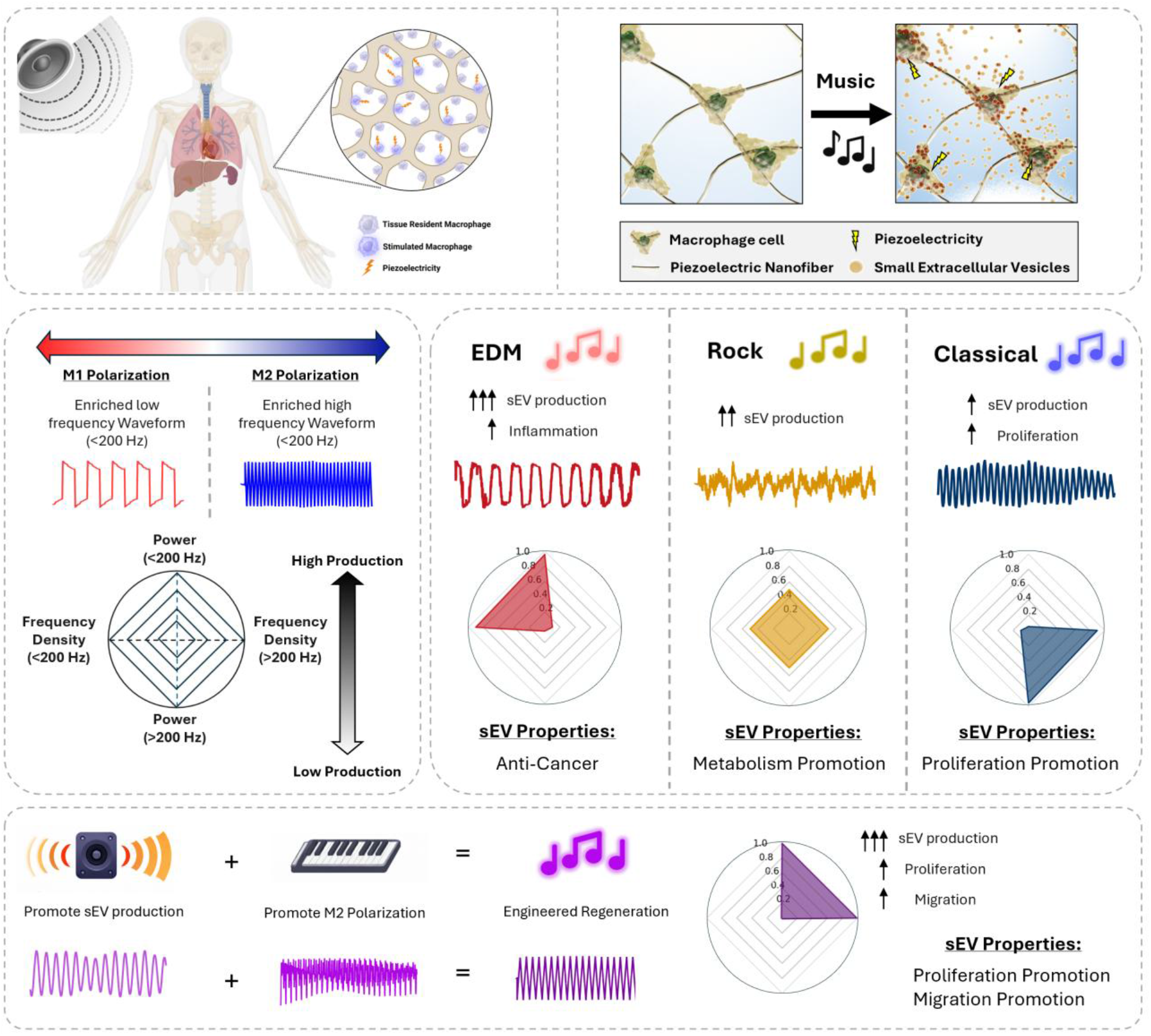
Overview schematic of the piezoelectric nanofiber substrate (PES) platform with various acoustic stimuli. The PES platform mimics the piezoelectrically sensitive extracellular matrix that macrophages adhere to in a tissue. Using this phenomenon, this platform can take advantage of acoustic frequency power and harmonic density to differentiate the small extracellular vesicle (sEV) production and cellular phenotype. EDM music utilizes low frequency power with high dissonance at low frequencies resulting in high sEV production, but with promoted inflammation. A selected classical music stimulus takes advantage of powerful high frequency, and high dissonance in frequencies above 200 Hz which promotes regeneration but suffers from low sEV Production. Custom-made regenerative music is a hybrid stimulus that takes advantage of both the power in low frequencies and the dissonance in high frequencies to promote sEV production while also promoting regeneration. Figure made in Biorender.

Human disease models have become increasingly important in drug development due to the high failure rate of drug candidates in clinical trials. Despite a 44% increase in R&D expenditures by the top 15 pharma companies since 2016, reaching $133 billion in 2021, the drug attrition rate hit an all-time high of 95% that year.^[1]^ Most drugs fail in clinical stages despite proven efficacy and safety in animal models, largely because clinical trial decisions rely almost exclusively on animal-derived data. This translational gap has led to no significant increase in new drug approvals despite growing investments in pharmaceutical R&D.^[2]^ While human disease models based on 2D culture have widely been used for high-content (HC) and high-throughput (HT) screening (HCHTS), 2D culture environments offer cells a relatively simple and uniform environment.^[3,4]^ Such models are not sufficient to represent complex phenomena, such as tissue structures,^[5,6]^ spatial cell populations,^[7,8]^ and dynamic physiological processes,^[9,10]^ leading to differences in drug deliveries and efficacies.^[11,12]^ Hence, there has been a critical need to develop more reliable and accurate human disease models *in vitro* to better mimic the native environment, including tissue architecture in 3D coordinates, cell-cell, cell-extracellular matrix (ECM), and ECM-drug interactions.^[13]^

To address the translational gap, 3D cell culture techniques have emerged as promising solutions, with various human disease models developed over recent decades, including cellular spheroids, organoids, and scaffolded tissue models.^[3]^ Cellular spheroids and organoids, as scaffold-free 3D micro-physiological systems, effectively replicate tissue structures and functions, creating a more physiologically relevant environment for cellular activities.^[9]^ By spontaneously mimicking *in vivo* conditions through the expression of ECM, they not only promote physiologically reliable cellular behavior (*e*.*g*., cell differentiation, proliferation, signaling, and drug response in *in-vivo* conditions)^[14,15]^ but also allow dynamically precise molecular behavior (*e*.*g*., diffusion, transport, accumulation, and drug delivery within the 3D cellular system).^[12,16]^ Furthermore, the spherical geometry of cellular spheroids offers a straightforward 3D spatial coordinate system that closely mimics real tissue, representing cell-cell, cell-ECM, or ECM-ECM interactions.^[17]^ This simplification facilitates efficient monitoring/tracking and mathematical modeling of drug transport and efficacies as well as cell responses within the 3D cellular environment.^[18–20]^ To achieve 3D HCHTS that are compatible with traditional drug screening methods^[3,21]^ and adaptable to advanced artificial intelligence (AI) or machine learning (ML) analysis,^[3,22]^ various methodologies, such as hanging-drop method,^[23]^ bioreactor culture,^[24]^ suspension culture,^[25]^ and well-plate-based pellet culture,^[26]^ have been introduced to parallelize the generation, culturing, and analysis of spheroids and organoids. However, these methods exhibit spheroid throughput/content scalability limitations and/or spheroid size dispersity,^[27]^ showing a notable trade-off between the two features. Additionally, many of these methodologies for 3D HCHTS require complicated operation or are not compatible with the conventional readouts for drug testing, further complicating their adoption in both static and dynamic systemic analysis (*e*.*g*., endpoint or one-point analysis; *in-situ*, continuous, or real-time analysis).^[3,28]^

## 2. Methods

### 2.1. Materials and Reagents

Polyacrylonitrile (PAN; 181315), sodium borohydride (213462), and Pluronic f-127 powder (9003-11-6) were purchased from Sigma Aldrich (MA). N,N-Dimethylformamide (DMF; D119-4) was purchased from Fischer Scientific (MA). Ethyl alcohol (3791-10), 99% acetic acid (BDH3092) and 0.22 μm vacuum filters (76010-388) were purchased from VWR (PA). Chitosan powder (c1569) was purchased from Spectrum Chemical (NJ). Minimum essential medium (MEM; 10-010-CV) and Corning SpinX centrifuge filters (431491) were purchased from Corning (NY). Fetal bovine serum (FBS; 26140079), 100× antibiotic–antimycotic (15240062), 0.25% trypsin–EDTA (25200072), and Prestoblue (A13261) were purchased from ThermoFisher (MA). 4% paraformaldehyde in 0.1 M phosphate buffer (15735) was purchased from Electron Microscopy Science (PA). Cell counting kit-8 (CCK-8; 850-039-kl01) was purchased from Enzo Life Sciences (NY). The LIVE/DEAD™ cell imaging kit (488/570) was purchased from Thermofisher Scientific (MA).

### 2.2. Preparation of PAN solution and PAN nanofiber scaffolds

Solution property characteristics were performed similarly to prior work. [12,45] Solution viscosity was measured using a CPA-40 spindle connected to a Brookfield DV-I Prime viscometer (Brookfield, Toronto, Canada). The rotational speed of the spindle was ramped up from 0.5 rpm to whichever speed at which the torque reached closest to 100% (at least above 95%). After confirming that viscosity was independent of the shear rate, the viscosity value at maximum torque was recorded. Surface tension was measured using an automatic surface tensiometer (QBZY-1; Shanghai Fangrui Instrument, Shanghai, China), which had a platinum-coated plate connected to a hook. The force exerted on the hook as the plate came in contact with the solution was converted into surface tension values Arduino code (Atlas Scientific, NY) was used to take electrical conductivity measurement through a glass-body electrical conductivity probe (K = 0.1, Oakton) paired with an embedded conductivity circuit (EZO-EC; Atlas Scientific, NY) and an Arduino Uno Rev3 board. All solution property measurements were taken at room temperature immediately before or after electrospinning to correlate them most closely with the resulting nanofiber properties. Nanofibers with a diameter of 760 nm were produced through an electrospinning process. A solution of 10 wt% PAN was prepared in DMF. Electrospinning was carried out under specific conditions, namely an electrospinning distance of 10 cm, an applied voltage of 13 kV, and a solution feed rate of 1 mL h^-1^. This process was conducted in a controlled environment of 23 °C and 40% relative humidity. The resulting nanofibers were collected on a rotating collector drum covered with aluminum foil, operating at 400 rpm. The electrospinning duration was optimized to achieve nanofibers with the desired thickness of approximately 100 µm.

### 2.3. Piezoelectric property measurement

Nanofiber scaffolds were prepared in a cantilever setup, similar to prior work. [12,45] This setup allows for the controlled application of strain to the samples while simultaneously measuring their electrical output. The PAN nanofiber scaffolds were cut into strips of size 4 × 1.2 cm, and brass slabs of size 7.2 × 1.6 × 0.01 cm, electrically isolated with polyimide tape, were employed as electrodes to measure the voltage. One brass slab was in direct contact with the nanofiber sample, secured with double-sided copper tape, while the other slab remained unexposed. Two 24-gauge wires were soldered to these electrodes, sealed with polyimide tape, and connected to a breadboard with inputs to a PicoScope 2204A (Pico Technology Ltd, Cambridgeshire, UK) for voltage measurement. To induce controlled strain, a 2.3 g proof mass was placed on the cantilever’s end, driven by a custom-made oscillatory system, with the cantilever holder clamped atop a subwoofer diaphragm. A sinusoidal sine wave with a controlled amplitude and frequency was applied to the speaker system, and the voltage output was measured.

### 2.4. Scanning electron microscopy images of PES

Scanning electron microscopy (SEM; Thermo Prima environmental-SEM; ThermoFisher, MA) was used to image samples with and without cells. Cell-free samples were fixed on a metallic stud (75210; Electron Microscopy Science, PA) with double sided conductive tape and sputter-coated with gold before imaging under 10 kV. The sample thickness, average fiber diameter, and pore size were measured based on the SEM images using the ImageJ software.

### 2.5. 2D Cell culture and media collection

Mouse macrophage cell line (RAW264.7) was purchased from the American Type Culture Collection (ATCC) and were cultured in T75 flasks at a density of 2E6 cells per well in DMEM/high glucose medium containing 10% FBS and 1% antibiotic-antimycotic. The medium was exchanged every 48 hours. Once the culture reached 70% confluency, the cells were washed with PBS 3 times, replaced with EV depleted medium, and incubated for 24 hours before collection for sEV isolation. Cells were then harvested after media collection with trypsin–EDTA for 5 min followed by spinning down the cells at 1100 rpm for 4 min. The cells were resuspended in fresh MEM and diluted appropriately to be counted using a hemocytometer.

### 2.6. PES Cell culture and media collection

The scaffolds were cut into 3 cm x 1.5 cm strips and processed for seeding, which included washing, coating with chitosan for cell adhesion, and sterilization. Each sample was rinsed in distilled water for 30 min and transferred to a solution of 1 mg mL−1 chitosan in 0.1 M acetic acid for 30 min. The coated nanofiber scaffolds were then rinsed with fresh distilled water for 30 min and air dried. The processed samples were placed in a 60 mm diameter Petri dish and UV-sterilized for 30 minutes before seeding. RAW264.7 cells were passaged and seeded separately at 5E5 cells per scaffold. The seeded scaffolds were then incubated for 30 min at 37 °C before the addition of serum-containing MEM and incubation at 37 °C and 5% CO2. After 48 h, the conditioned medium was replaced with serum-free medium and incubated for 24 h before being collected for sEV isolation. Cells were then counted after media collection using a cell counting kit-8 assay: The culture medium was aspirated, and the scaffolds were washed with PBS 3 times before adding 3% CCK-8 reagent in the cell culture media and incubation for 2 hours at 37 °C and 5% CO2. The cell count was then measured through the absorbance at 460 nm wavelength.

### 2.7. Cell seeding efficiency for PES culture

A cell seeding efficiency measurement testing was performed based on a previous study (ref): RAW264.7 cells were seeded separately on 2 mm × 2 mm scaffolds (*N* = 3) with and without a chitosan coating at a density of 5 × 10^3 cells per scaffold and incubated in serum-containing MEM at 37 °C and 5% CO2 for 24 h. The cultured medium was aspirated, and the scaffolds were washed with PBS thrice before the addition of 10% CCK-8 reagent in cell culture media and incubation for 4 h at 37 °C and 5% CO2. The cell count was then measured through absorbance at 460 nm wavelength. Chitosan coated scaffolds were used for future cell culture tests.

### 2.8. Cell proliferation test

RAW264.7 cells were seeded separately on 2 mm × 2 mm scaffolds (*N* = 3) at 1E3 cells per scaffold and incubated at different set time periods (0.5, 1, 2, 3, 5, 7, 9, 11, 13 days). Media was replaced every day, and confluency was monitored. The media was aspirated, and the scaffolds were washed with PBS thrice before the addition of 10% CCK-8 reagent in cell culture media and incubation for 4 h at 37 °C and 5% CO2. The cell count was then measured through absorbance at 460 nm wavelength.

### 2.9. Acoustic environment characterization

The acoustic stimulation environment was characterized using a microphone coupled with the DecibelX application. The volume and pitch settings were adjusted to achieve an appropriate average amplitude and frequency according to the DecibelX application. The average amplitude and frequency was measured over a period of 5 minutes in an acoustic controlled environment blocking external noise. Acoustic environments were created with a range of 50-110 dB amplitude and 50-1000 Hz frequency using this process.

### 2.10. sEV isolation and concentration measurement

sEVs were isolated from the media *via* size-based separation. The isolated media first went through a 0.22 μm filter to capture larger vesicles and cell debris. The flowthrough solution was then added to a 100 kDa centrifuge filter and centrifuged at 200*g* for 4 × 30 min time periods; the mixture was washed with PBS between each centrifuge session. The sEV solution was then concentrated down to 1 mL for sEV characterization. The sEV concentration and size distribution were measured through nanoparticle tracking analysis (NTA; Nanosight NS300; Malvern, Worcestershire, UK). Samples were diluted appropriately to maintain accurate particle counts. For each sample, five 60 s videos were acquired at a camera level of 8 and detection threshold of 2. The laser chamber was cleaned with milliQ water between each sample reading to ensure no sample contamination occurred. The videos were analyzed using the NTA3.0 software to obtain particle concentration, along with the mean and mode particle sizes of each sample.

### 2.11. Acoustic stimulation and media collection

PES cultures containing serum-free medium were stimulated using sinusoidal acoustic waves (3-Ω subwoofer; PS-EW1-2; Samsung Electronics, Suwon-si, Republic of Korea) in an acoustically controlled box. Samples were stimulated for 15 minutes at various amplitude levels (50-110 dB) and frequencies (50-1000 Hz) levels. The samples were incubated at 37 °C and 5% CO2 for 24 hours before being collected for sEV isolation. For the music study, the PES cultures were stimulated with the appropriate song for 15 minutes followed by incubation at 37 °C and 5% CO2 for 24 hours before being collected for sEV isolation.

### 2.12. Acoustic characterization of Music Tracks

An audio input file was imported into a python code and the time-based frequency, waveform, and overall frequencies were plotted. Waveform was determined through plotting the wave amplitude as a function of time. The frequency was determined through applying a fast Fourier transform on the wave amplitude. Stimuli plotted in the study included basic sine waveforms at different frequencies, square waveforms, triangle waveforms, sawtooth waveforms, instruments played at 250 Hz Frequency, and tested music tracks.

### 2.13. Dissonance calculations of Music Tracks

Dissonance *D*_*f*_ was calculated and summed up over each frequency at a given time using the following equations:

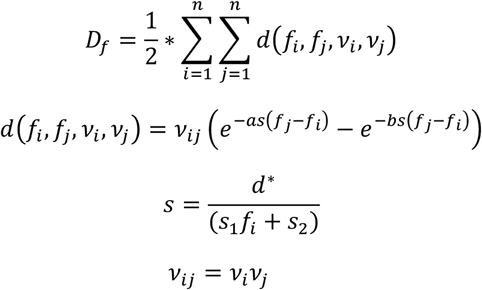

Where *f* is frequency, *v* is amplitude, and s is an interpolation parameter between two frequencies which are determined to be 0.021 and 19 based on least squared fit modeling. The constants a and b, and d* were deciphered constants from previous calculations of local dissonance. [46] The average dissonance was calculated over the time-length of the stimuli and normalized between 0 and 1. Consonance was then calculated using the following equation:

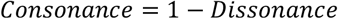

These values were used as the horizontal portion of the radar plot.

### 2.14. Cytokine release measurement of cells on stimulated PES

IL6 and IL10 cytokines were detected using an ELISA kit from Thermofisher (IL6 – 88-7064-88, IL10 - 88-7105-88). The manufacturer’s protocol was followed for the test: 96 well ELISA plates were coated with capture antibody in the coating buffer. The plates were sealed and incubated overnight at 4C. The wells were aspirated and washed 3 times with PBS. Each sample was soaked for 1 minute for each wash step followed by blotting the plate on absorbent paper to remove any residual buffer. Wells were then blocked with ELISA Diluent (1X) and incubated at room temperature for 1 hour. The diluent was aspirated and washed once followed by the incubation of antigen standard and culture supernatant overnight at 4C. The sample was aspirated and washed 3 times followed by adding the antibody detection to all wells and incubating at room temperature for 1 hour. The wells were aspirated and washed 3 more times. Avidin-HRP (IL6) or Streptavidin-HRP (IL10) was added and incubated at room temperature for 1 hour, followed by 5 more washes. TMB solution was added to each well and incubated at room temperature for 15 minutes. Stop solution (2M sulfuric acid) was added to each well and the absorbance at 450 nm was measured using a microplate reader. Protein concentration was calculated using the standard curve absorbance measurements.

### 2.15. Flow Cytometry

Cells were detached using enzyme-free dissociation buffer and the stained with either CD86-PE, or CD206-PE for 30 minutes at 4C. Cells were washed 5 times with FACS buffer and tested for fluorescence using a cytek Norther Lights (NL)-CLC flow cytometer. Positive controls for CD86 consisted of RAW264.7 cells that were stimulated with LPS for 12 hours before detachment. Positive controls for CD206 consisted of RAW264.7 cells that were stimulated with IL10 for 12 hours before detachment. Fluorescence results were analyzed through FlowJo.

### 2.16. Western blot

sEVs were lysed with 1× RIPA buffer (9806; Cell Signaling Technology, USA), and the total protein concentration was quantified using pierce BCA protein assay kits (23225; Thermo Fisher, USA). The protein amount in lysed sEVs was estimated based on a calibration curve plotted by albumin (BSA) standards. Next, 12 μg of proteins from sEV lysates were denatured and loaded on sodium dodecyl-sulfate polyacrylamide gel electrophoresis (SDS-PAGE). The separated proteins were subsequently transferred onto a nitrocellulose membrane (1662807; Bio-Rad, USA) and blotted overnight with primary antibodies purchased from Santa Cruz Biotechnology, USA: anti-CD9 antibody (C-4), anti-CD63 antibody (MX-49.129.5), anti-CD47 antibody (B-11), anti-ALIX antibody, anti-TSG101 antibody (W27), The secondary antibodies (anti-mouse HRP-linked antibody, 7076; Cell Signaling Technology, USA) were then treated for blotting and the HRP on the immunoblots was detected by Clarity Max Western Enhanced Chemiluminescence (ECL) substrate (1705060; Bio-Rad, USA) using a ChemiDoc XRS+ system (Bio-Rad, USA).

### 2.17. Transmission electron microscopy of sEVs

The sEV solutions were negatively stained and imaged through transmission electron microscopy (TEM) using the Talios F200i (S)TEM (ThermoFisher, MA) at an 80 kV accelerating voltage. The TEM samples were prepared by adding 3 μL of 1 × 10^9 particles per mL sEV solution to an ultrathin carbon film copper grid and incubating at room temperature for 2 min. The solution was then aspirated using filter paper and washed with 3 μL filtered distilled water for 10 s. After aspirating the distilled water, 3 μL of Uranyless negative staining solution (22409; Electron Microscopy Science, PA) was added to the sample grid (CF200-CU-25; Electron Microscopy Science, PA) and incubated for 1 min. The Uranyless solution was aspirated, and the grid was left to dry for 2 minutes before imaging. The sEV particle diameter was measured using the default distance measurement tool in ImageJ.

### 2.18. RNA sequencing and analysis

RNA was isolated from both RAW264.7 cells and sEVs using the Qiagen miRNeasy Tissue/Cells Advanced Kit using the manufacturer’s protocol. Buffer AL was added to a pellet of cells or 700 ul sEV solution at 10^10^ sEV/ml and incubated at room temperature for 3 minutes followed by transferring the lysate to a gDNA eliminator spin column and centrifuged for 30 seconds at 8000 x g. The flow through was saved and 1.3X volume of isopropanol was added. The mixture was transferred to a RNeasy UCP MinElute spin column and centrifuged for 15 s at 8000 g, discarding the flowthrough. Buffer RWT was added and centrifuged through, discarding the flowthrough. Buffer RPE was added and centrifuged through, discarding the flowthrough. 80% ethanol was added and centrifuged, discarding the flowthrough. RNase-free water was added directly to the center of the spin column membrane and incubated for 1 minute before centrifuging for 1 minute at 12000 g. The RNA solution was tested for purity using a nanodrop absorbance spectrometer before RNA sequencing. RNA sequencing was performed through a third-party company (HaploX, Hong Kong). Gene quantification was performed using HTSeq, and differential sequencing was conducted between each sample using DESeq2. Functional enrichment analysis was performed using the Enrichr software tool. [47–49]

### 2.19. MS-based proteomics of sEVs

sEVs were lysed in 1X RIPA buffer. A total of 100 ug of protein from exosomes were trypsinized and eluted using an S-Trap™ Micro Spin Column. The eluted protein was then dried down and resuspended in a mobile phase buffer. The suspension was loaded into an ultrahigh-resolution mass spectrometer coupled to a nano-ultrahigh-pressure liquid chromatography for bottom-up proteomics.

### 2.20. sEV efficacy test on breast cancer cells

A CCK-8 assay used to short term viability of sEVs from different music genres were tested on MCF-7 cells purchased from the American Type Culture Collection (ATCC). MCF-7 cells were seeded into a transparent 96-well plate at a concentration of 1E4 cells/well. After 24 hours, sEVs were added and transfected for 48 hours at 37C. 10% CCK-8 reagent was added, and the cells were incubated for 4 hours at 37C. Cell viability was measured by absorbance at 460 nm and was normalized to the control group without sEVs. For long-term effects, a clonogenic assay was conducted. MCF-7 cells were seeded into transparent 6-well plates at a seeding density of 4E4 cell/well. After 24 hours, sEVs were added and transfected for 48 hours at 37C. Cell were then washed, trypsinized, and reseeded into 6 well plates at 500 cells/well. Medium was replaced every 48 hours. After 1 week, the medium was aspirated and the cells were washed with PBS three times. Colonies were fixed with 6% glutaraldehyde for 5 minutes at room temperature and stained with 0.5% (W/V) crystal violet for 15 minutes at room temperature. Each well was photographed, and colony numbers were quantified using ImageJ. The relative colony formation (%) was normalized by the control group without sEVs.

### 2.21. Testing efficacy of sEVs on the cell proliferation of MCF7 cells

MCF7 cells were seeded into a transparent 96 well plate and incubated at different set time periods (0.5, 1, 2, 3, 5, 7, 9, 11, 13 days). Media supplemented with sEVs at 1E9 EV/well was replaced every day, and confluency was monitored. The media was aspirated, and the scaffolds were washed with PBS thrice before the addition of 10% CCK-8 reagent in cell culture media and incubation for 4 h at 37 °C and 5% CO2. The cell count was then measured through absorbance at 460 nm wavelength.

### 2.22. Testing the effect of sEVs on MCF7 migration (scratch assay)

MCF7 cells were incubated with sEVs for 48 hours on a 6 well plate. Sterilized 20 ul pipet tips were used to make scratches. The scratched gaps were monitored at 0, 12, and 24 hours after the scratch under a light optical microscope. The closure of the gaps was quantified using ImageJ. The relatively scratched area was normalized to the control group.

### 2.23. Statistical analysis

The data presented in this paper are expressed as the mean

± the standard deviation (S.D.) or the standard error of replicate measurements (S.E.M.) and analyzed using one way ANOVA test using GraphPad Prism 9 software, where applicable.

## 3. Results

### 3.1. Design of PES, and acoustic-based optimization of sEV production

We successfully synthesized the PES, (**Fig. 2A-B, and Fig. S1**) exhibiting robust piezoelectric transduction, converting acoustic-driven mechanical stress into a measurable electrical output in PBS (**Fig. 2C**). The piezoelectric output of PES can be tuned by adjusting amplitude and frequency of the acoustic input (**Fig. 2D**). The amplitude of the acoustic input was directly proportional to the peak-to-peak output, plateauing at 191.7 ± 14.6 mV (**Fig. 2E**). In contrast, the frequency of the input was inversely proportional to the piezoelectric output, with a maximum voltage of 311.5 +/-15.5 mV (**Fig. 2F**). Varying the waveform shape of the acoustic input can alter the shape of the electric output, while possessing similar correlations between frequency and amplitude and electric output as the basic sine wave input (**Fig. S2**). As a control, the piezoelectric activity of PES can be reduced by up to ∼90% through post-synthesis thermal treatment, providing an equivalent substrate for mechanistic studies (**Fig. S3**). To improve cell adhesion and biocompatibility of the PES, we coated chitosan on the material, resulting in a 91.2 ± 10.8% viability of RAW264.7 (**Fig. S4**). The piezo response of the chitosan coated PES remains within a suitable range to stimulate the cell membrane and generated a similar pattern to pristine PES for further studies.

**Figure 2.**
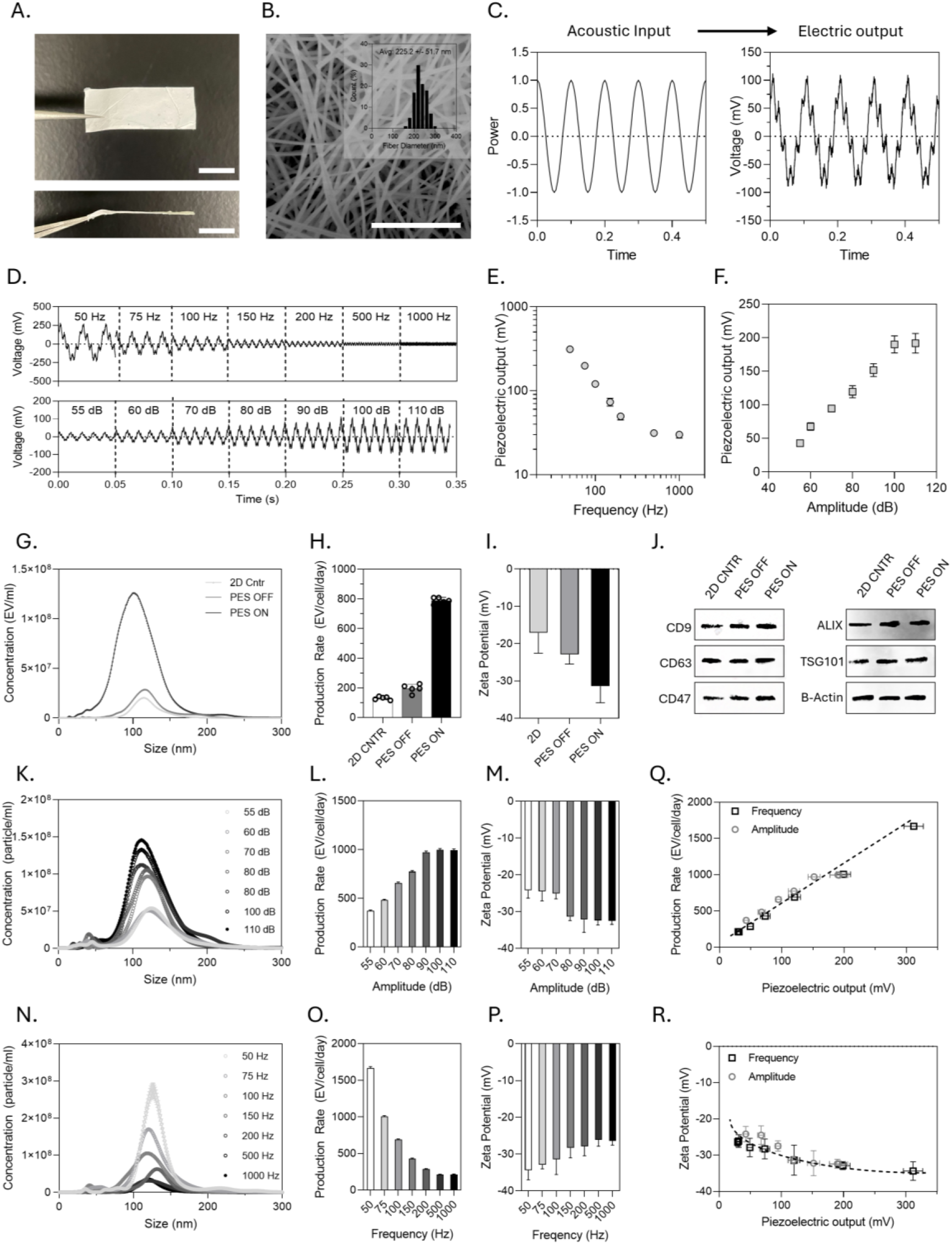
Characterization of the Piezoelectric substrate (PES) and evaluation of sEV production. **A.** Photographic representation of the PES mat. Scale bar is 1 cm **B**. Scanning electron micrograph of the microstructure of the PES mat. (inset) The average fiber diameter of the fiber-mat used in future experiments are 225 +/-51 nm. Scale bar is 5 µm. **C**. Graphical representation of the piezoelectric effect converting acoustic mechanical forces to electrical output for piezoelectric stimulation. **D**. The relationship of acoustic parameters in wave frequency and amplitude on the piezo-responsive behavior of the fiber-mat. Overall, the frequency is inversely related to the peak-to-peak output (**E**), and the amplitude is directly proportional to the peak-to-peak output (**F**). **G-J**. Production calculations and characterization of the RAW264.7 derived sEVs from various culturing conditions including standard monolayer 2D culture (2D Cntr), PES platform without acoustic stimulation (PES OFF), and the PES platform with acoustic stimulation. **G**. Size distribution and concentration calculations of sEVs derived from each condition using nanoparticle tracking analysis (NTA). **H**. Production rate calculation measured through concentration with NTA, and normalized by the culturing time, and cell count through manual cell counting. Units of production rate are EV/cell/day. **I**. Zeta potential measurements of sEVs derived from each condition using DLS. **J**. Validation of the production of sEVs through sEV-derived protein marker characterization using western blot. **K-M**. Relationship between amplitude of acoustic input on **K**. size distribution, **L** sEV production rate, and **M**. zeta potential. **N-P**. Relationship between Frequency of acoustic input on **N**. size distribution, **O** sEV production rate, and **P** zeta potential. **Q**. Linear correlation between sEV production and the piezoelectric output. **R**. Inverse correlation between zeta potential and the piezoelectric output.

We measured and compared the sEV production rate of RAW264.7 cells in traditional 2D culture, PES without stimulation (PES OFF), and PES with acoustic stimuli (PES ON) under 100 Hz, and 80 dB acoustic waves (**Fig.2G-H, and Fig. S5**). PES ON group successfully produced sEVs 6.7-fold faster than 2D culture control, and 4.1-fold faster than PES OFF group, indicating that the PES stimulated by acoustics promoted sEV secretion. To note, deactivated PES with a stimulation reduced to 19.44 ± 1.99 mV led to a minimal increase in production rate (1.6-fold) compared to 2D culture, thus the change was solely due to the piezo effect rather than acoustics. PES ON derived sEVs showed the highest charges compared to the other sEVs (**Fig. 2I**), which indicated good dispersity in physiological solutions. Positive western blot signals for sEV surface markers (CD9, CD63, CD47), and cargo (ALIX, TSG101) confirmed the abundant existence of sEVs in all groups (**Fig. 2J and Fig.S6**). Lastly, transmission electron microscopy (TEM) imaging of all samples (**Fig.S7**) showed round morphology with similar size distributions to nanoparticle tracking analysis (NTA) data (**Fig.2G**). We also successfully tuned the sEV production rate through adjusting the acoustic parameters on the same PES material. We observed a linear increase in sEV production rate when increasing the amplitude of stimuli, whereas the acoustic frequency had an inverse relationship with sEV production rate (**Fig. 2K-P**). Interestingly, this trend was observed universally when we adjusted the waveform of the stimuli (**Fig. S8**). When comparing sEV production rate with piezoelectric output, we saw a linearly positive trend with sEV production rate, suggesting the piezoelectric output was a driving force in sEV production (**Fig. 2Q**). This was supported by observing a linear decrease in sEV production when stimulated with gradually heat deactivated PES material under the same acoustic stimuli (**Fig.S3**). Interestingly, when comparing zeta potential with piezoelectric effect, we observed an inverse relationship (**Fig. 2R**). Overall, stimulated PES promoted sEV production from RAW264.7 cells by as high as 15.5-fold (50 Hz, 80 dB stimuli) compared to monolayer 2D culture systems and this sEV production can be tuned through controlling acoustic parameters and thus piezoelectric stimuli.

### 3.2. Music-inspired control of sEV production and cell programming

We used music assemblies spanning common listening genres (e.g. EDM, classical, rock) as standardized audible acoustic waves to investigate how musically distinct stimuli influence sEV production and macrophage phenotypes. The music assembles chosen for each genre were (1) *On the floor ultrafunk* by KPHK to represent EDM, (2) *Don’t Go Breaking My Heart* by Elton John and Kiki Dee to represent rock, and Bach’s *Suite in B minor, BWV1067: V. Polonaise* by Johann Sebastian to represent classical music. To understand how the acoustic features differentially engage piezoelectric substrates and cellular responses, we extracted each track’s waveform, time-resolved spectral content, bulk frequency distribution, and acoustic dissonance (**Fig. 3A**). These three music assemblies possessed significant differences in sound pressure (power), spectral composition (frequency), and harmonic dissonance. First, the EDM assembly possessed strong energy concentrated in bass and sub-bass bands (20-200 Hz) at high sound pressures. In addition to the powerful sub bass frequencies, EDM displayed near-maximal normalized dissonance (0.98 ± 0.05) (see **Methods**), resulting in considerable amplitude instability, previously reported to promote systemic proinflammatory responses under short term exposures (**Fig. 3A.i**). [28–30] In contrast, the classical assembly was characterized by powerful signals at high frequencies, outside the primary piezoelectric activation range of the PES. Classical music also exhibited minimal dissonance (0.02 ± 0.01), describing a stable, harmonious construction of frequencies, previously reported to promote systemic anti-inflammation outcome (**Fig. 3A.ii**). [20,21] The rock assembly fell between these extremes, with broader spectral balance and intermediate dissonance (0.51 ± 0.05) (**Fig. 3A.iii**). When comparing bulk frequency information with dissonance calculations, we observed a consistent trend: high signal density occurring at sub-bass frequencies was a driving force in promoting acoustic dissonance. To test whether this relationship generalizes beyond music, we performed parallel analyses using canonical waveforms (triangle, square, and sawtooth). As expected, waveforms with higher harmonic density exhibited higher dissonance, with sawtooth > square > triangle (**Fig. S9-11**).

**Figure 3.**
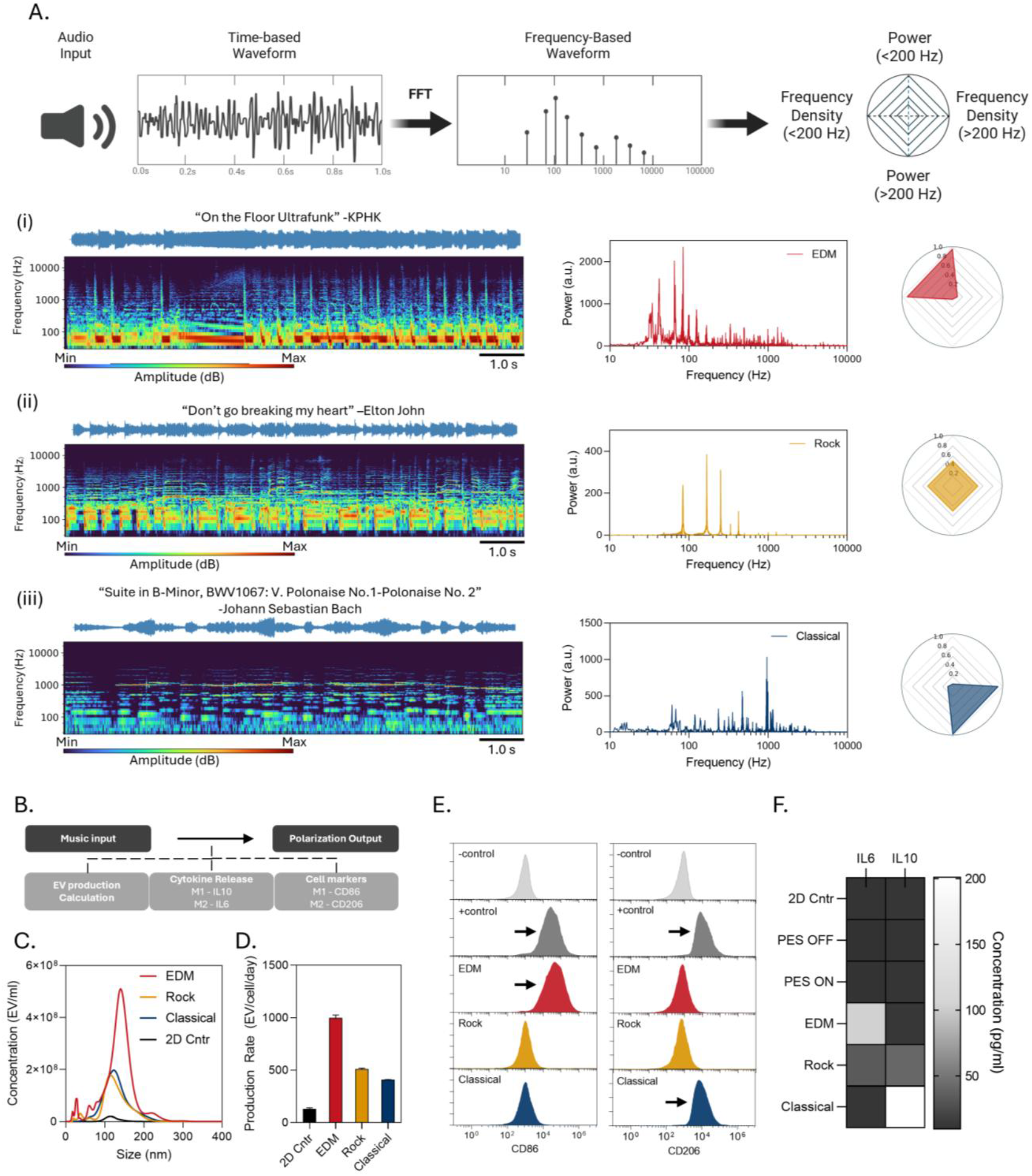
Music-inspired control of sEV production and macrophage phenotype. A Schematic representation of the acoustic characterization process starting with an audio input, and obtaining the time-based waveform, the fast Fourier transform frequency-based information, and a radar plot characterization the (top) total acoustic power at frequencies below 200 Hz, (left) Frequency density at frequencies lower than 200 Hz, (bottom) Overall power at frequencies above 200 Hz, and (right) frequency density at frequencies above 200 Hz. Acoustic characterization was performed for 3 musical samples. (i) An electronic music sample containing powerful low frequency waves, and high density at <100 Hz frequencies, resulting in a radar plot at the top-left position. (ii) A rock music sample containing a balance both power and density in low and high frequencies resulting in a radar plot at the center. (iii) A classical music sample containing a powerful high frequency wave, and high density in >200 Hz frequencies, resulting in a radar plot at the bottom-right position. **B-F**. Testing the effects of the acoustic stimuli input on the sEV production and macrophage phenotype of RAW264.7 cells through sEV production calculation, cytokine release measurements, and Cell marker characterization. **B**. A flow chart representation of the characterization process to observe the immune response and sEV production output based on a given music input. **C-D**. sEV production measurements of the 3 musical stimuli. **E**. Surface marker characterization of CD86 (M1 polarization) and CD206 (M2 polarization). Negative controls and the positive control samples were measured to compare the signal with the musical stimuli. **F**. Measurement of cytokine concentration released into the cell culture supernatant of the control groups, and the various musical stimuli groups. Cytokine concentrations were measured using ELISA (N=5, mean ± s.d).

To understand how music-driven activation of the PES influences sEV biogenesis and macrophage polarization, RAW264.7 cells were cultured on PES and applied defined acoustic stimuli (waveforms and music assemblies) and then quantified functional outputs at the vesicle and cellular levels. Specifically, we measured (i) sEV production rate, (ii) polarization-associated cytokine release (IL-6 and IL-10), and (iii) macrophage surface marker expression (CD86 and CD206) (**Fig. 3B-D**). As a result, all music stimuli promoted sEV production compared to 2D culture system (**Fig. 3E**).

EDM promoted the highest sEV production enhancement (991.5 ± 21.4 EV/cell/day) followed by Rock (502.7 ± 6.7 EV/cell/day) and classical (429.4 ± 7.9 EV/cell/day) (**Fig. 3B-D**). Interestingly, the power at low frequencies (<200 Hz) followed the sEV production trend (EDM > rock > classical) (**Fig. 3A)**. EDM’s stronger power at low frequencies increased piezoelectric stimulation, inducing greater sEV production. [13] When we assessed macrophage polarization under music stimuli, highly dissonant EDM stimuli (Dissonance = 0.98 ± 0.05) induced a significant increase in CD86 expression and IL6 cytokine release, suggesting M1 polarization of macrophages (**Fig. 3E-F**). In contrast, highly consonant classical music stimuli (dissonant = 0.02 ± 0.01) promotes CD206 expression and IL10 cytokine release, suggesting M2 polarization (**Fig. 3E-F**). Lastly, we observed no significant change in surface markers or cytokine release under a balanced dissonant rock stimulus (dissonance = 0.51 ± 0.05) (**Fig. 3E-F**). To support this correlation between dissonance and polarization, we compared different levels of dissonant waveforms (sawtooth > square > triangle) as controls and found highly dissonant sawtooth (0.86 ± 0.12), and square waves (0.74 ± 0.22) promoted M1 polarization, while low dissonant triangle waves (0.24 ± 0.19) promoted M2 polarization (**Fig. S12-15**). We confirmed the morphological stability and size distribution of sEVs from all groups through TEM and NTA (**Fig. S16-17**). Overall, these results showed that macrophage polarization can be controlled by tuning the dissonance of the acoustic stimuli, highlighting how musically distinct inputs, and the piezoelectric simulation they elicit, can program macrophage state.

### 3.3. Cargo analysis and *In Vitro* application of reprogrammed sEVs

We next performed RNA sequencing and proteomics profiling to validate music-driven cell phenotype shifts and determine whether they were reflected in sEVs. Across all conditions, the global RNA signatures of sEVs broadly tracked those of their parental cells, consistent with phenotype-dependent vesicle biogenesis. Differential expression analysis revealed robust, stimulus-specific divergence across music conditions (**Fig. 4A**). Pathway enrichment analysis further highlighted distinct biological programs. In EDM-stimulated cells, adaptive immune processes and inflammatory pathways were significantly upregulated, whereas gene sets associated with cell cycle progression, cell division, and proliferation were downregulated. In contrast, classical stimulation produced the opposite trend, with enrichment of proliferation-associated pathways and suppression of inflammation signaling. Rock-stimulated cells possessed similar expressions to the 2D control, exhibiting modest changes across these pathways (**Fig. 4B and Fig. S18**). Under all music conditions, the expression of CACNA (voltage-gated Ca^2+^ channel) gene family were upregulated relative to the 2D controls, and positively correlated with sEV production rate, suggesting Ca^2+^ influx as a driver of enhanced vesicle release (**Fig. 4C**). We next profiled sEV proteomic cargos and surface markers across music-stimulated conditions and benchmarked them against 2D culture and PES OFF controls (**Fig. S19**). All groups exhibited sEV surface markers (CD63, CD9, CD47) and cargos (ALIX, TSG101). Consistent with the observed cellular polarization, EDM-sEVs possessed M1 cargos in the form of proinflammatory cytokines (IL6, IL7, IL16), immune activation markers (CD86, CD84, CD27, etc.), and tumor growth-inhibitor proteins (SCAI, MIIP). In contrast, classical-sEVs possessed M2 cargos in the form of anti-inflammatory markers (CD206), matrix metalloproteinases (MMP3, MMP13) and collagen subunits (COL27A1, COL18A1, COL11A2) for bone and cartilage regeneration. To further investigate the potential therapeutic implications, we investigated the intrinsic therapeutic effects of EDM-sEVs and classical-sEVs *in vitro*. Interestingly, EDM-sEVs promoted cytotoxicity in breast cancer cells compared to the 2D-control-sEVs, with a decrease in cell density (**Fig. 4D-F**). We also observed inhibition of cell proliferation and thus insignificant gap closure from breast cancer cells incubated with EDM-sEVs, further supporting their anti-cancer potential (**Fig. 4G-I**). In contrast, classical-sEV promoted cell growth as observed in the enhanced cell density compared to 2D-control-sEVs (**Fig. 4D-F**), while accelerating gap closure, and long-term proliferation, demonstrating regeneration potentials (**Fig. 4G-I, and Fig. S20**).

**Figure 4.**
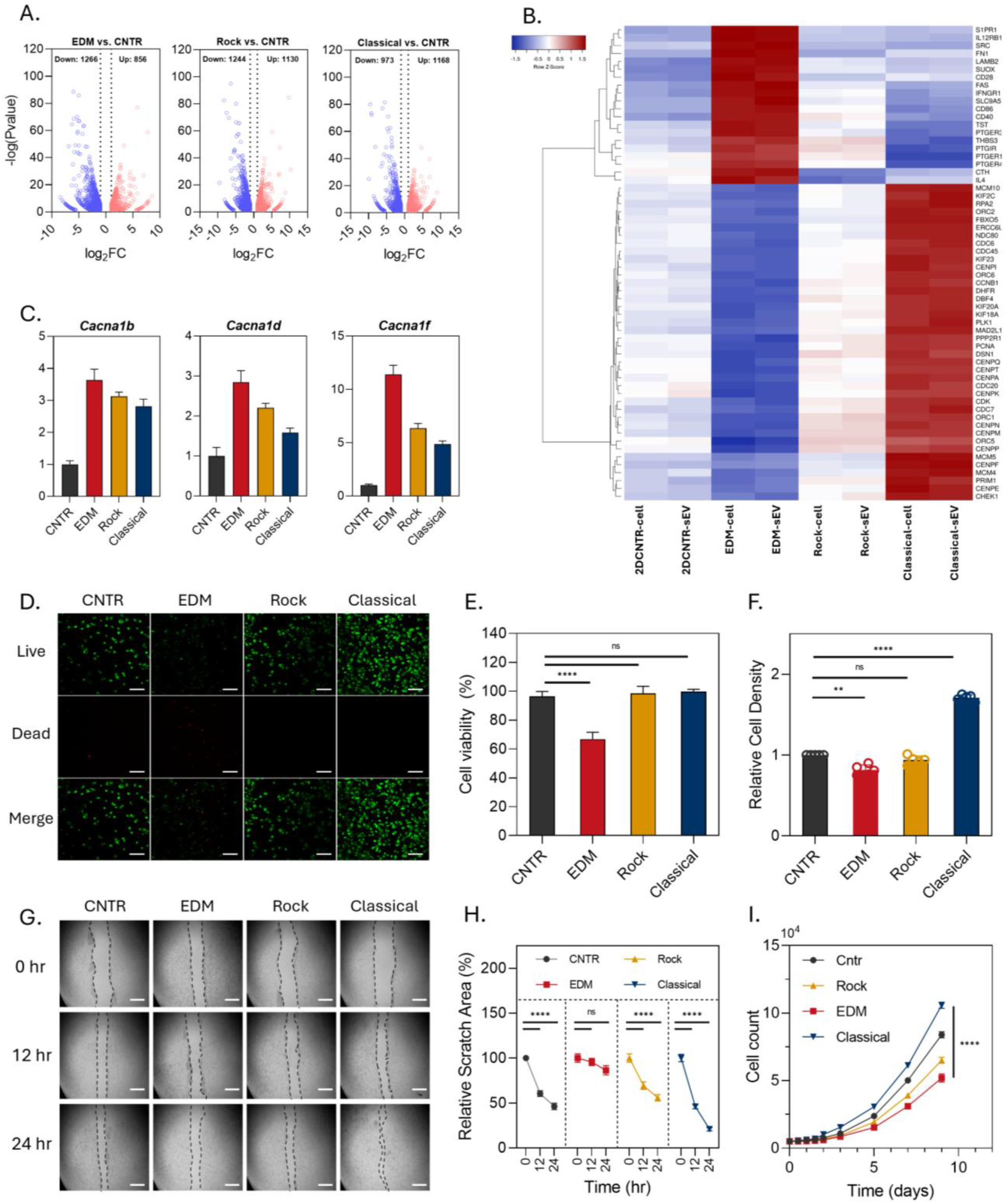
Cellular and sEV RNA and function vary with music stimuli. **A**. Differential sequencing between EDM, Rock, and Classical music stimuli compared to the 2D controlled condition. **B**. Clustered heatmap quantifying expression correlated to inflammation (top) and cell growth (bottom) **C**. Relative expression of CACNA1 family RNA responsible for voltage-gated calcium channel protein expression. **D-F** characterization of anti-cancer properties of music derived sEVs when incubated with MCF7 cells **D**. measurements of the natural sEV toxicity to MCF7 cells using a Live/Dead assay. Scale bar is 50um. **E**. calculated cell viability from the live/dead assay. (N=3, mean ± s.d.) **F**. calculated cell density **G-H** characterization of regenerative properties of music derived sEVs when incubated with hFOB1.19 cells. **G**. Scratch assay with 1000 EVs/cell dosage of sEV. **H**. Quantitative measurements of the scratch area (N=5, mean ± s.d). **I**. long-term proliferation study characterizing sEVs effect on MCF7 cell proliferation. (N=3, mean ± s.d) One-way ANOVA with Tukey’s post-test. Ns: not significant; *p>0.05; **p<0.01; ***p<0.001; ****p<0.0001.

### 3.4. Engineering regenerative stimuli for enhanced production of anti-inflammatory sEVs

To integrate the high yield sEV production elicited by EDM with the pro-regenerative bioactivity associated with the classical stimuli, we designed a composite music assembly (“Engineered Regeneration”) that combined the key acoustic elements underlying each effect. The assembly incorporated (i) a sub-frequency bass within the resonance frequency of PES to promote sEV production, (ii) a piano chord progression that promotes M2 polarization suitable for regeneration applications, and (iii) water sound as a negative control that does not affect the cells. Piano was chosen after screening 13 instrument samples, where it most consistently promoted M2 polarization due to its low dissonance timbre (**Fig. S21-24**). Acoustic analysis confirmed that the Engineered Regeneration music assembly combined high power within the PES-optimal frequency window, driven by the bass component (peak-to-peak voltage = 211.12 ± 18.44 mV), with a harmonically stable, low-dissonance musical structure provided by the piano chords (**Fig. 5A, and Fig. S25**). We then deconvoluted the contributions of each module by stimulating RAW264.7 cells with the individual components and defined combinations, and quantifying both vesicle output and polarization markers. Stimuli that contain the bass component (**Fig. 5 B-D**: 1, 4, 6, 7) promote sEV production up to 8.2-fold higher than 2D culture, indicating sub-bass–driven piezoelectric activation as the dominant determinant of vesicle yield (**Fig. 5B and Fig. S26**). In parallel, stimuli containing the piano component (**Fig. 5 B-D:** 2, 4, 5, 7) promoted M2 polarization characterized by CD206 expression and IL10 release, identifying that the consonant piano progression is sufficient to promote an M2-like program under these conditions (**Fig. 5C-D**). Across all engineered conditions, sEVs maintained round morphology and comparable size distribution (**Fig. S27-28**), supporting that the stimulus primarily reprogrammed output and bioactivity rather than gross vesicle structure. Finally, we evaluated the functional consequence of the engineered regimen in a regenerative context. Engineered-Regeneration-sEVs increased hFOB1.19 osteoblast cell density, ∼3-fold enhancements relative to 3D control sEVs and accelerated scratch closure ∼2-fold faster, achieving 100% closure within 24 hours (**Fig. 5E-G**). These results demonstrated a rational, modular approach to “composing” acoustic–piezoelectric stimuli that jointly maximize sEV manufacturing and program therapeutic function, highlighting the PES platform as a tunable route to application-specific, phenotype-imprinted vesicle therapies.

**Figure 5.**
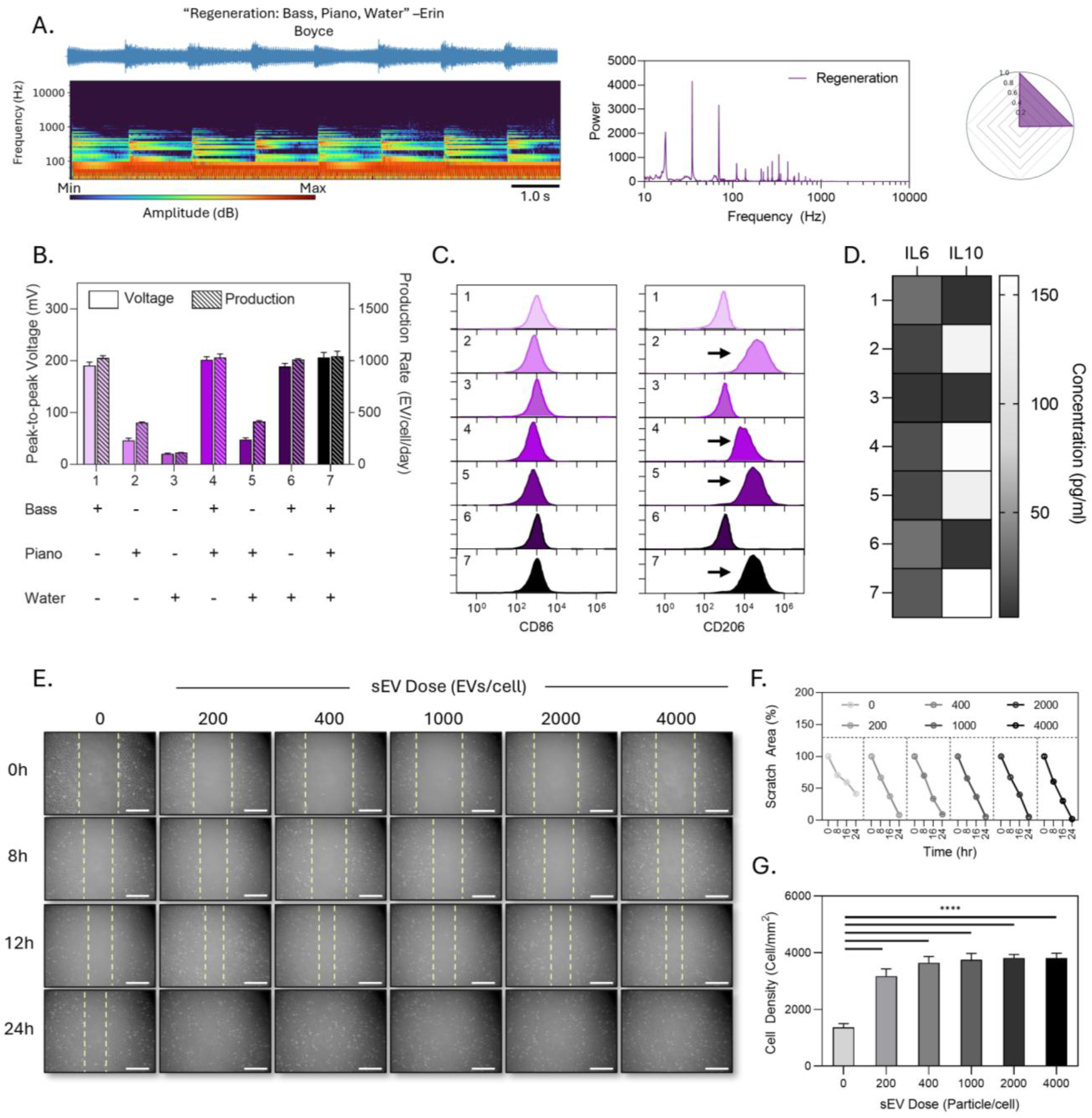
Characterization of custom music optimized for promoting sEV production and M2 polarization for regeneration potentials. **A**. acoustic characterization of the regeneration music. **B**. A component analysis showing the impact of each acoustic component on piezoresponsive behavior (solid) and sEV production rate (striped), (N=5, mean ± s.d.) **C**. FACS characterization of CD86 and CD206 markers of each component. **D**. IL6 and IL10 cytokine measurements through ELISA assays (N=5, mean). **(E-G)** Characterization of tissue regeneration potential of regeneration music derived sEVs on hFOB1.19 cells. **E**. Scratch assay with a range of the sEV dosages. **F**. Quantification of the scratch area with respect to time (N=3, mean ± s.d.) **G**. Cell density measurements of hFOB1.19 after incubation of regeneration music derived sEVs. (N=3, mean ± s.d.) Quantitative measurements of the scratch area (N=5, mean ± s.d). One-way ANOVA with Tukey’s post-test. Ns: not significant; *p>0.05; **p<0.01; ***p<0.001; ****p<0.0001.

## 4. Discussion

In this study, we leveraged our PES platform to connect audible acoustic inputs to bioelectrical cell stimulation and thereby control three coupled outputs: (1) sEV production rate, (2) macrophage phenotype and the resulting sEV cargo, and (3) the intrinsic therapeutic activity of the produced sEVs *in vitro*. Building on these, we further demonstrated a rational “compositional” strategy, an engineered music stimulus, that simultaneously maximizes vesicle yield and tunes vesicle function toward tissue regeneration, providing a tunable route to fast, controlled production of phenotype-imprinted sEVs.

A key mechanistic question is what drives the increase in sEV output under PES stimulation. Prior approaches typically enhance vesicle production using either high-voltage electrical stimulation to trigger Ca^2+^ influx or high-frequency mechanical stimulation (e.g., surface acoustic waves) to engage mechanotransduction [15,31–33]. In our platform, acoustic excitation generates both mechanical vibration and piezoelectric charge; however, two observations point to piezoelectricity as the dominant contributor. First, thermal deactivation of PES piezoelectricity produced a proportional, linear reduction in sEV production rate, implicating the electrical transduction pathway as necessary for the yield gains (**Fig. S3**). Second, the low-frequency regime that most strongly boosted production (<200 Hz) is unlikely to deliver the MHz–GHz mechanical cues typically required to enhance vesicle release through purely mechanical pathways [33]. Thus, PES enables a sound-to-electric route for scalable sEV production in an easily programmable frequency band. Importantly, this platform adds a capability largely absent from existing stimulation-based manufacturing: by shaping the acoustic “timbre” (including harmonic structure/dissonance), we can tune macrophage polarization and thereby tune the functional subtype of macrophage-derived sEVs illustrated by the engineered regenerative stimulus (**Fig. 5**).

Multi-omic profiling further indicates that music-dependent shifts in cell state are imprinted onto vesicle composition. EDM stimulation produced a proinflammatory, immune-activation–enriched cellular program, supported by sEV cargo enriched for inflammatory cytokines and activation markers, [34–36] together with proteins linked to suppression of tumor growth and invasion (e.g., SCAI, MIIP). [37–40] These signatures align with the observed anti-proliferative and anti-migratory activity of EDM-sEVs *in vitro*. In contrast, classical stimulation drives cells towards proliferation-associated programs while dampening inflammatory pathways; correspondingly, classical-sEVs promoted proliferation and migration and were enriched for ECM remodeling and osteochondral-associated proteins (including MMPs and collagen subunits), consistent with regenerative function. [41–44] Overall, these data support a model in which fundamental acoustic structure, transduced through PES, acts upstream of macrophage polarization, which then determines sEV bioactivity.

This work also has practical implications for sEV biomanufacturing. The ability to independently and modularly control yield and function suggest a generalizable framework for designing acoustic–piezoelectric regimens matched to therapeutic intent without genetic engineering or extensive biochemical conditioning. Meanwhile, several limitations remain. Although production is accelerated, the downstream workflow remains similar to standard adherent culture and would benefit from integration with automated, closed-loop collection and purification to reduce labor and improve throughput. In addition, the current implementation favors adherent cells; extending these concepts to suspension systems will be important to broaden compatibility to other clinically relevant producers. Addressing these engineering constraints should further position music-guided acoustic–piezoelectric stimulation as a scalable and programmable route to therapeutic sEVs.

## 5. Conclusion

Our work establishes a programmable, noninvasive strategy to jointly control sEV yield and function through precisely defined acoustic “music assemblies.” By tuning frequency and amplitude, we significantly increased sEV production while simultaneously directing sEV bioactivities. Importantly, music assemblies revealed a mechanistic link between waveform characteristics and macrophage fate: dissonant, powerful low-frequency stimuli biased RAW264.7 cells toward an M1-like inflammatory state, whereas consonant higher-frequency stimuli promoted M2-like polarization and a regenerative milieu. These polarization shifts were reflected in sEV cargos, generating vesicles with distinct, cell-mimetic bioactivities. Leveraging this insight, we engineered a custom stimulus that maximized both sEV output and M2 polarization, producing vesicles that enhanced osteoblast growth. Together, these findings position PES as a tunable, universal piezoresponsive platform that bridges acoustic-based interventions with sEV engineering, enabling scalable, cargo-tailored, and application-ready sEV therapeutics.

## CRediT authorship contribution statement

**James Johnston:** Conceptualization, Methodology, Investigation, Formal Analysis, Visualization, Writing – Original Draft. **Erin Boyce:** Conceptualization, Methodology. **Tiago Thomaz Migliati Zanon:** Investigation **Hyunsu Jeon:** Visualization **Courtney Khong:** Investigation **Yun Young Choi:** Investigation **Nosang Vincent Myung:** Methodology **Martin Nunez:** Investigation **Mae-Lin Pinkstaff:** Investigation **Yichun Wang:** Conceptualization, Funding Acquisition, Supervision, Writing – Original Draft.

## Acknowledgements

This work was supported by the National Institute of Health (R35GM150608, and R21CA277663-01A1) and the Department of Education (P200A210048). We acknowledge the Analytical Science and Engineering Core Facility (ASEND) and Dr. Karl Cronberger for their technical assistance in PES fabrication and characterization. We acknowledge the Harper Cancer Research Institute Tissue Bank Facility (HCRI tissue-bank) and Dr. Emily Cronberger for their technical assistance with sEV particle analysis and Flow cytometry. We acknowledge the Notre Dame Integrated Imaging Facility and Dr. Sara Cole, and Dr. Maksym Zhukovskyi for their technical assistance with TEM, and the confocal microscopy. We acknowledge the Notre Dame Mass Spectrometry & Proteomics Facility, along with Dr. Bill Boggess, for their technical assistance with sEV proteomics. We acknowledge HaploX for their technical assistance with RNA-sequencing and analysis. We thank Gaeun Kim, Tiger Shi, Dr. Yichen Liu, Dr. Wei Zhang, Farbod Shirinichi, and Yao Huo for their technical and conceptual guidance.

## Data Availability Statement

The data that support the findings of this study are available from the corresponding author upon reasonable request.

