## Supplementary material for "Music-Inspired Acoustic-Piezoelectric Stimulation Accelerates Extracellular Vesicle Production and Programs Therapeutic Function": SI figures

[2]Independent Music Producer

**Supplemental figures:**

A.

| Pan wt% | Viscosity (cP) | Surface Tension (dynes/cm) | Nanofiber Diameter (nm) |
|---|---|---|---|
| 4 | 68.1 | 26.2 | No fiber formed |
| 6 | 98.4 | 26.4 | 225 +/- 51 |
| 8 | 192.8 | 27.3 | 350 +/- 77 |
| 10 | 335.4 | 26.4 | 476 +/- 103 |
| 11 | 419.5 | 26.8 | 1011 +/- 145 |
| 13 | 689.4 | 27.4 | 2205 +/- 521 |

B.

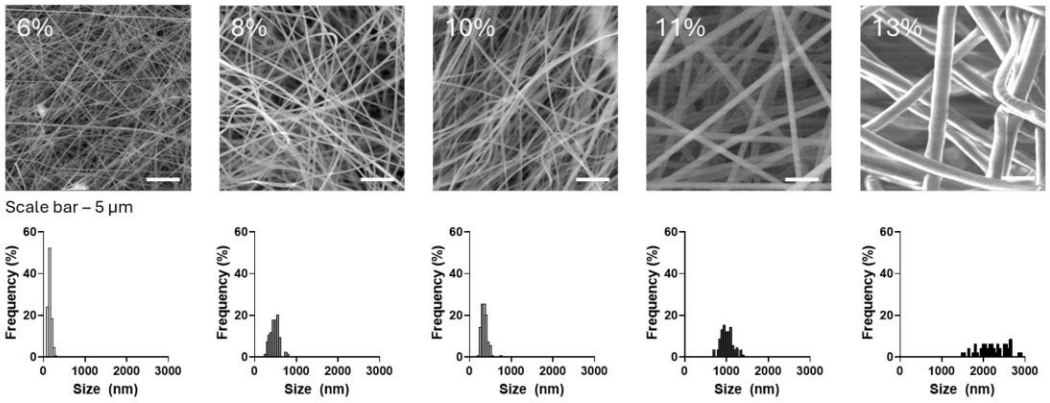


**Figure S1**: Optimization of the PAN concentration to control the fiber diameter. A. Viscosity, surface tension and nanofiber diameter measurements were conducted with varying PAN concentrations, and B. the fiber diameter distribution was plotted from SEM imaging. Scale bar is 5 um. (N=5)

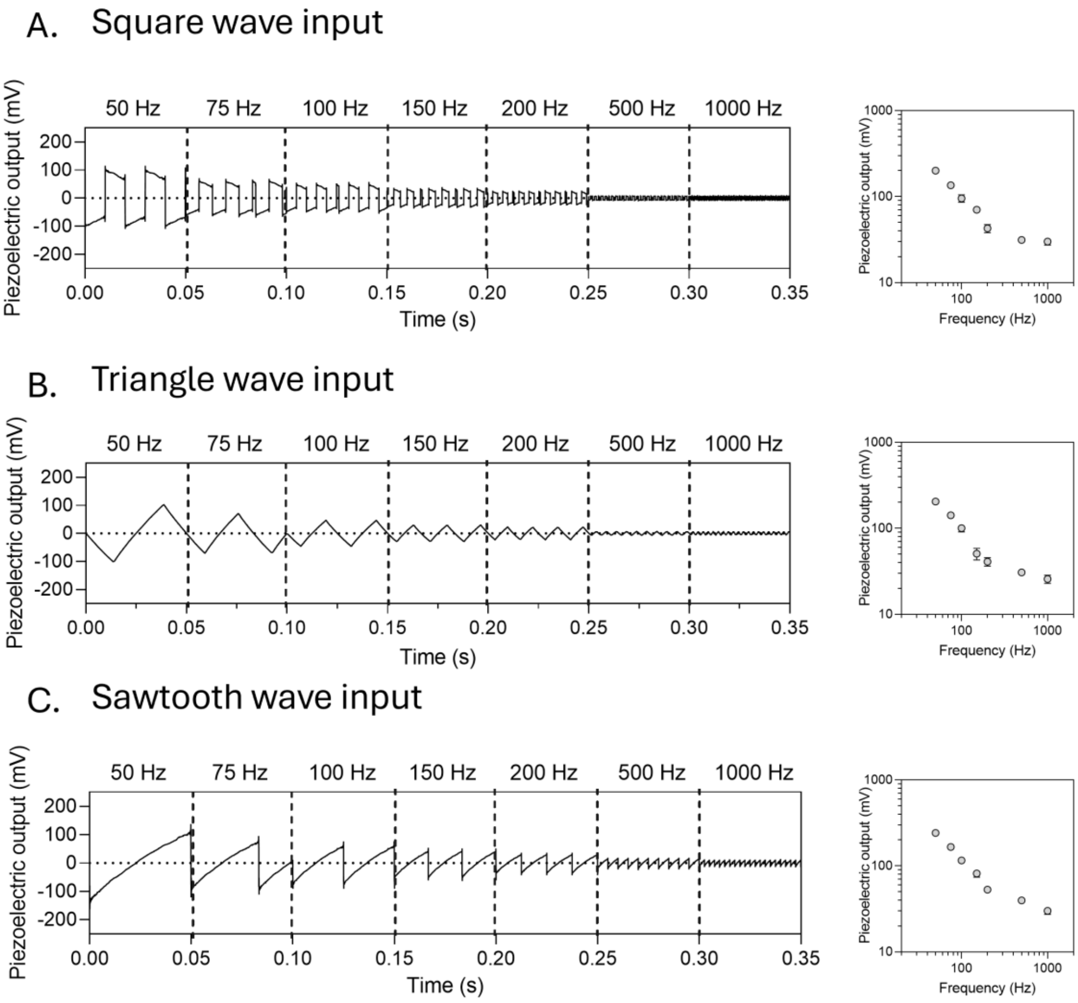


**Figure S2**: The effect of frequency and different waveforms on the piezoresponsive profiles of the PES material through a cantilever test. Piezoresponsive testing was conducted at various frequencies between 50-1000 Hz of A. Square waves, B. Triangle Waves, and C. Sawtooth waves.

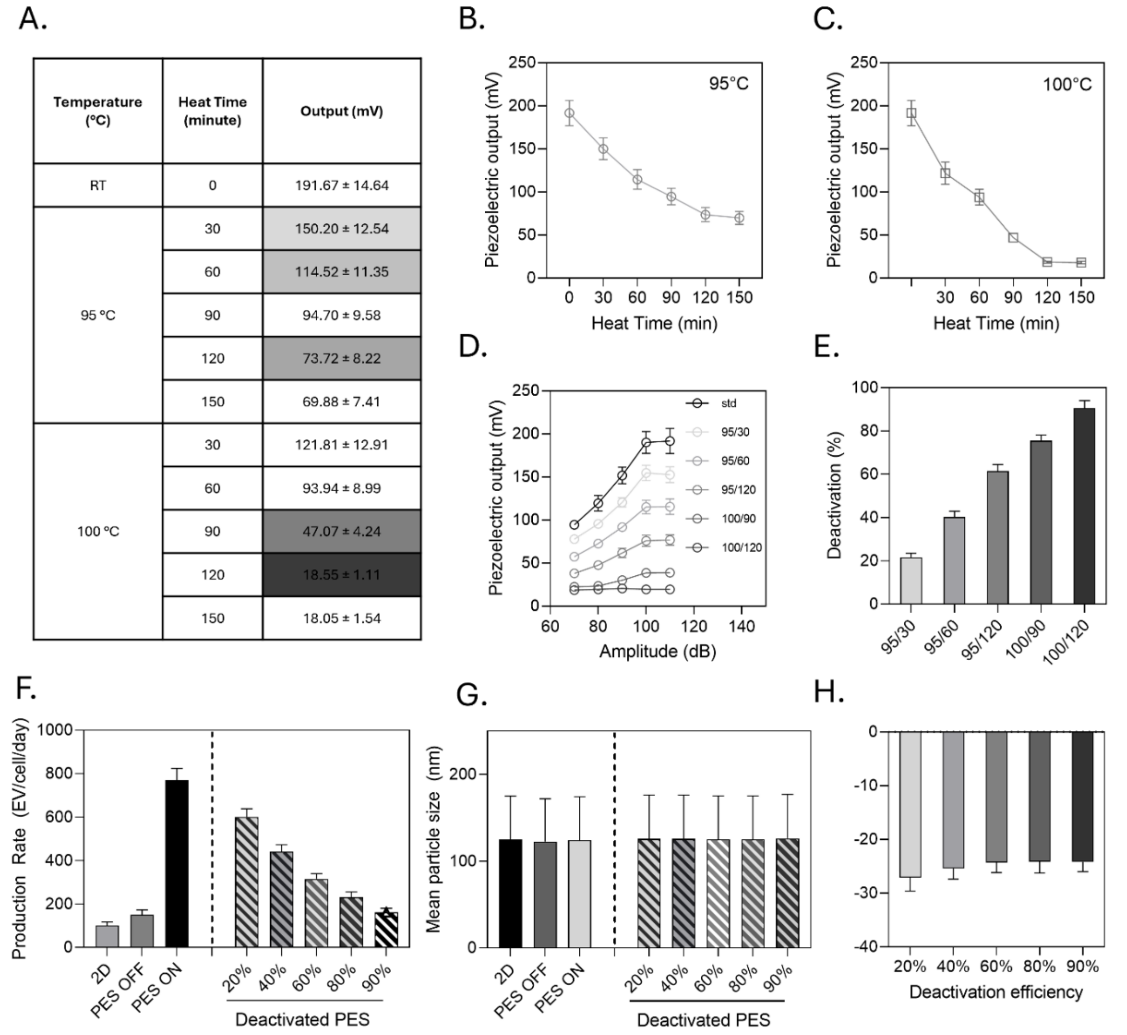

| Temperature (°C) | Heat Time (minute) | Output (mV) |
|---|---|---|
| RT | 0 | 191.67 ± 14.64 |
| 95 °C | 30 | 150.20 ± 12.54 |
| | 60 | 114.52 ± 11.35 |
| | 90 | 94.70 ± 9.58 |
| | 120 | 73.72 ± 8.22 |
| | 150 | 69.88 ± 7.41 |
| 100 °C | 30 | 121.81 ± 12.91 |
| | 60 | 93.94 ± 8.99 |
| | 90 | 47.07 ± 4.24 |
| | 120 | 18.55 ± 1.11 |
| | 150 | 18.05 ± 1.54 |



**Figure S3**: Effect of the crystalline-based piezoresponsive behavior on sEV production rate using one singular acoustic stimulus. (Sine wave, 100 Hz, 80 dB). A. A table of various temperatures and annealing times with measured piezoresponsive outputs (N=5). B. Effect of heat time at 95C on peak-to-peak piezoresponsive behavior. C. Effect of heat time at 100C on peakto peak piezoresponsive behavior. D. further amplitude studies on piezoresponsive behavior using optimized annealing conditions. E. Effect of optimized annealing conditions on peak-to-peak output. F. Effect of annealing on sEV production compared to the standard samples previously measured. G. Mean particle size of sEVs produced, and H. Mean zeta potential of sEVs produced.

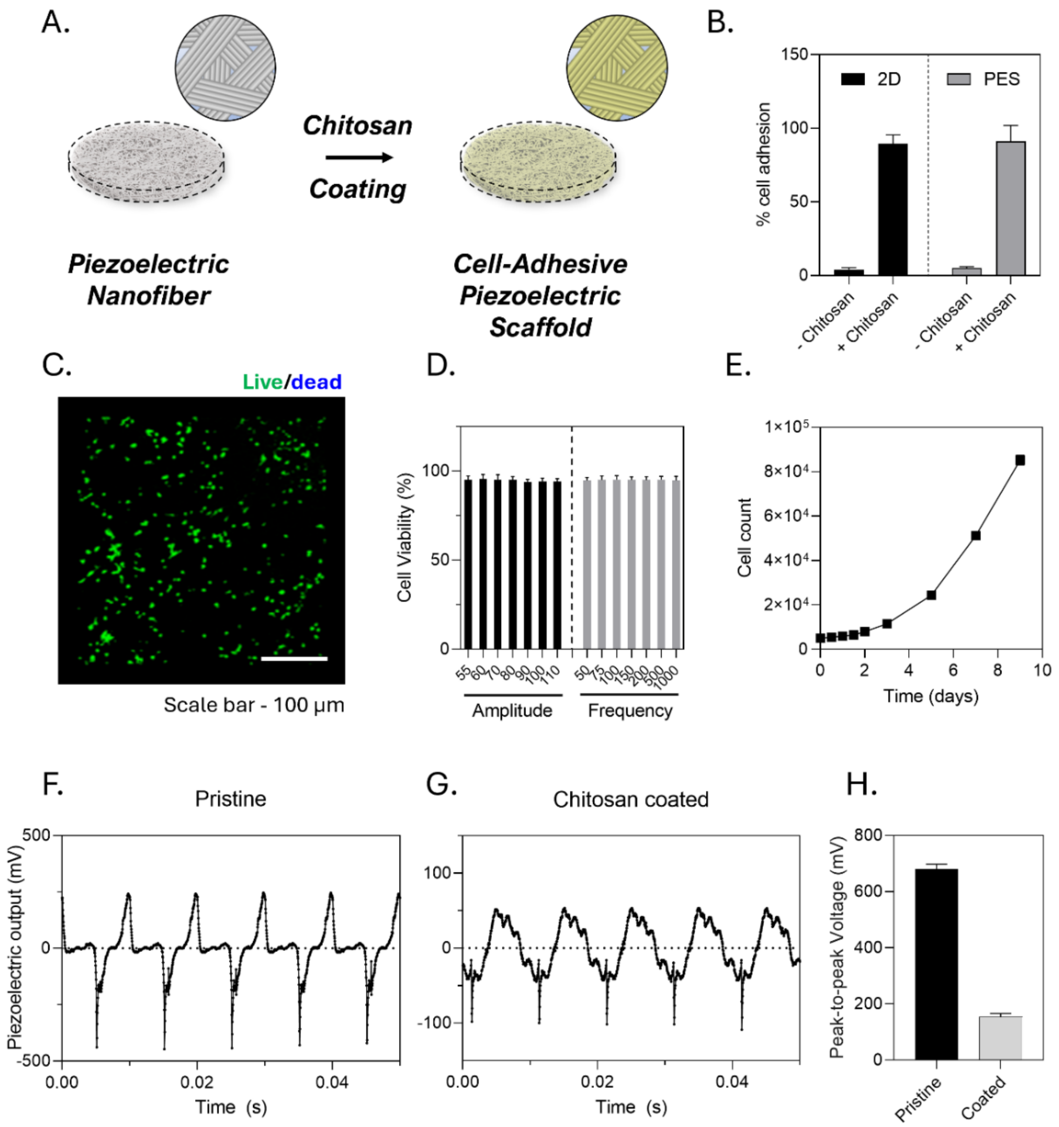


**Figure S4**: Cell culture characterization of the PES platform. A. Schematic of the chitosan coating process to enhance cell adhesive properties of the PES material. B. Cell adhesion testing using Cell counting Kit before and after washing (N=5). C. Confocal Microscopy Z-stacks of RAW264.7 cells on the PES material. D. Cell viability of RAW264.7 cells after various amplitudes and frequency of PES ON stimulation. Viability was calculated using CCK8 assay. E. Cell proliferation assay conducted using CCK8 assay over 9 days. F-H. Piezoelectric output of F. Pristine as-spun PES, and G. dry PES after chitosan coating using 100 Hz, 80 dB acoustic input. H. quantitative analysis of the peak-to-peak voltage.

A.

$$Rate\ (EV cell^{-1} day^{-1}) = \frac{EV\ yield}{Cell\ count * Incubation\ time}$$

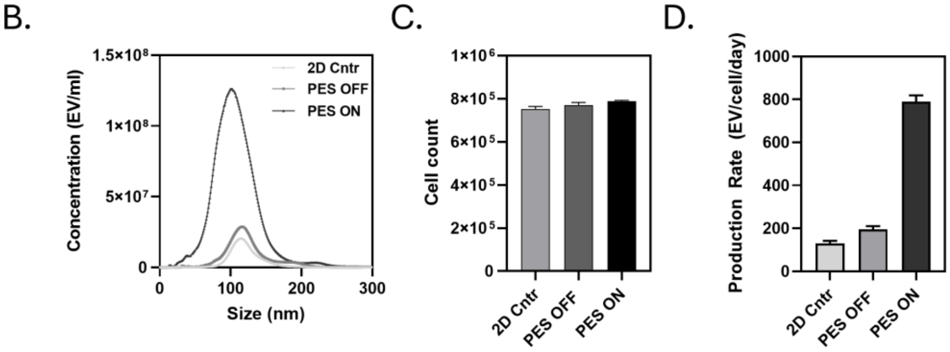


**Figure S5**: sEV production calculation process and sample calculation of production rate for RAW 264.7 cells under 2D CNTR, PES OFF, and PES ON conditions. A. The equation for sEV production rate being related to the EV yield, cell count, and incubation time for a universal productivity calculation. B. EV concentration and EV yield measurements provided through nanoparticle tracking analysis (NTA) of 2D CNTR, PES OFF, and PES ON conditions (N=5). C. Cell count measurements through manual cell counting after detachment of the cells from the PES material (N=5). D. Calculated production rate of sEVs from EV yield, and cell count data.

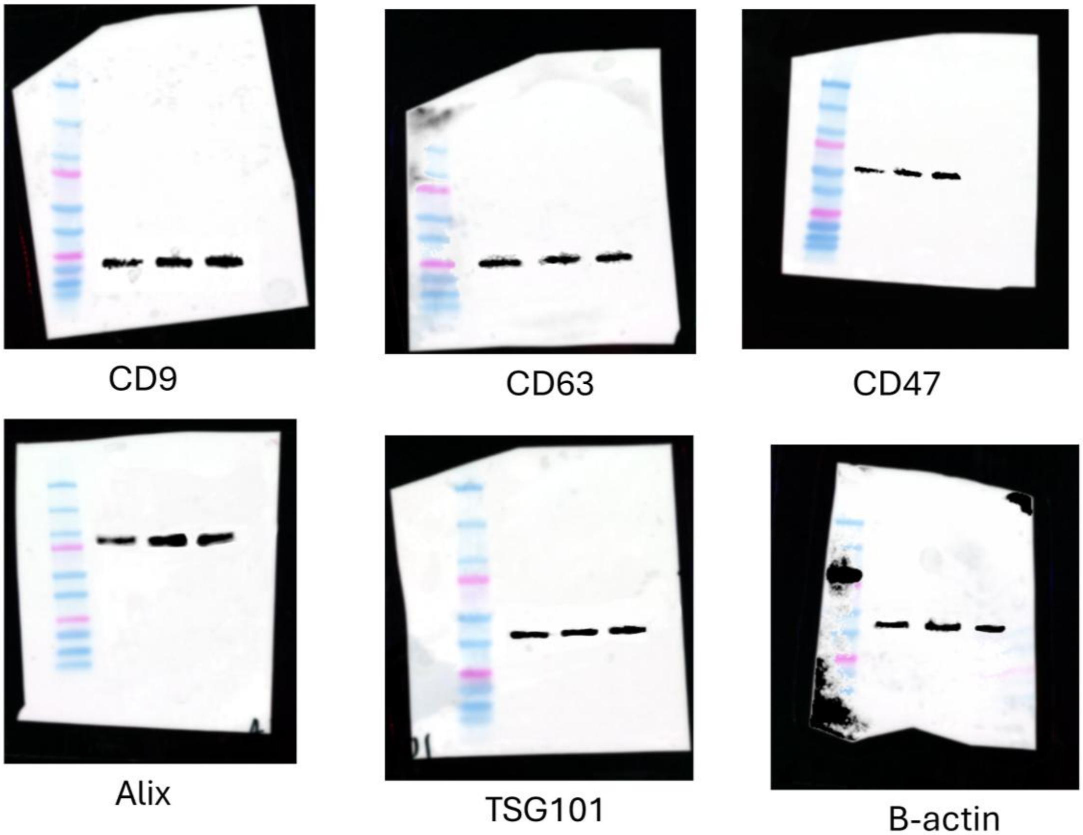


**Figure S6**: Raw Western blot gel images showing sEV marker characterization from RAW264.7 cells in 2D cntr, PES OFF, and PES ON conditions.

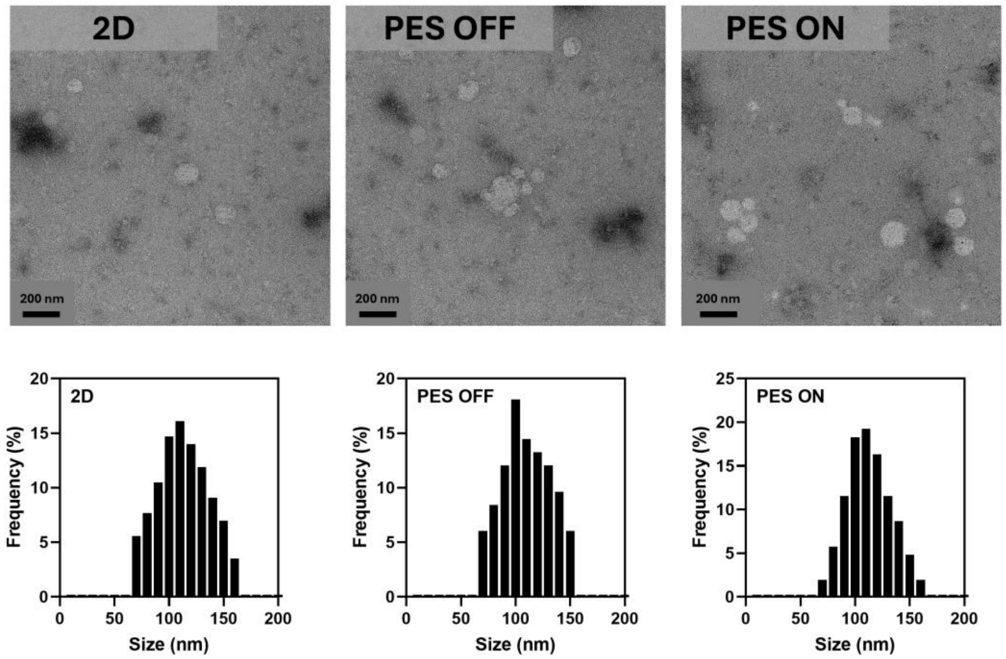


**Figure S7**: TEM imaging of sEVs produced from 2D, PES OFF, and PES ON samples with size distribution analysis (100 particles analyzed)

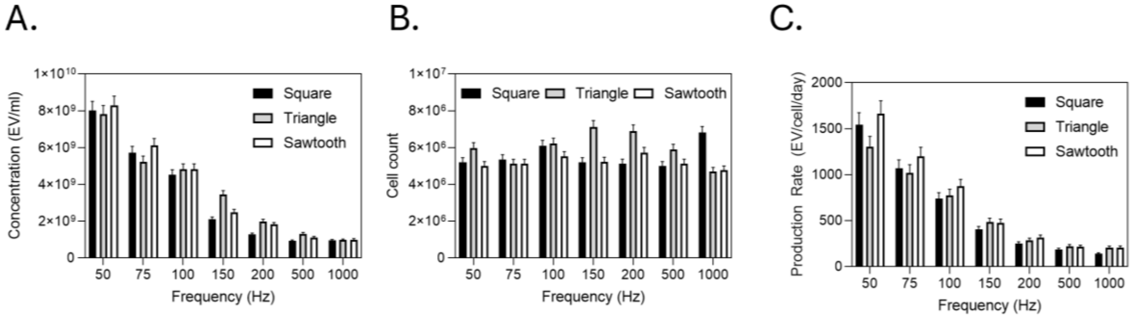


**Figure S8**: sEV production testing and verification of the effect on waveform input on sEV production at various frequency inputs. A. Concentration measurements using NTA (N=5). B. Cell count measurements using manual cell counting after detaching the cells from the PES material (N=5). C. Production rate calculation from the previous measurements at 24-hour incubation.

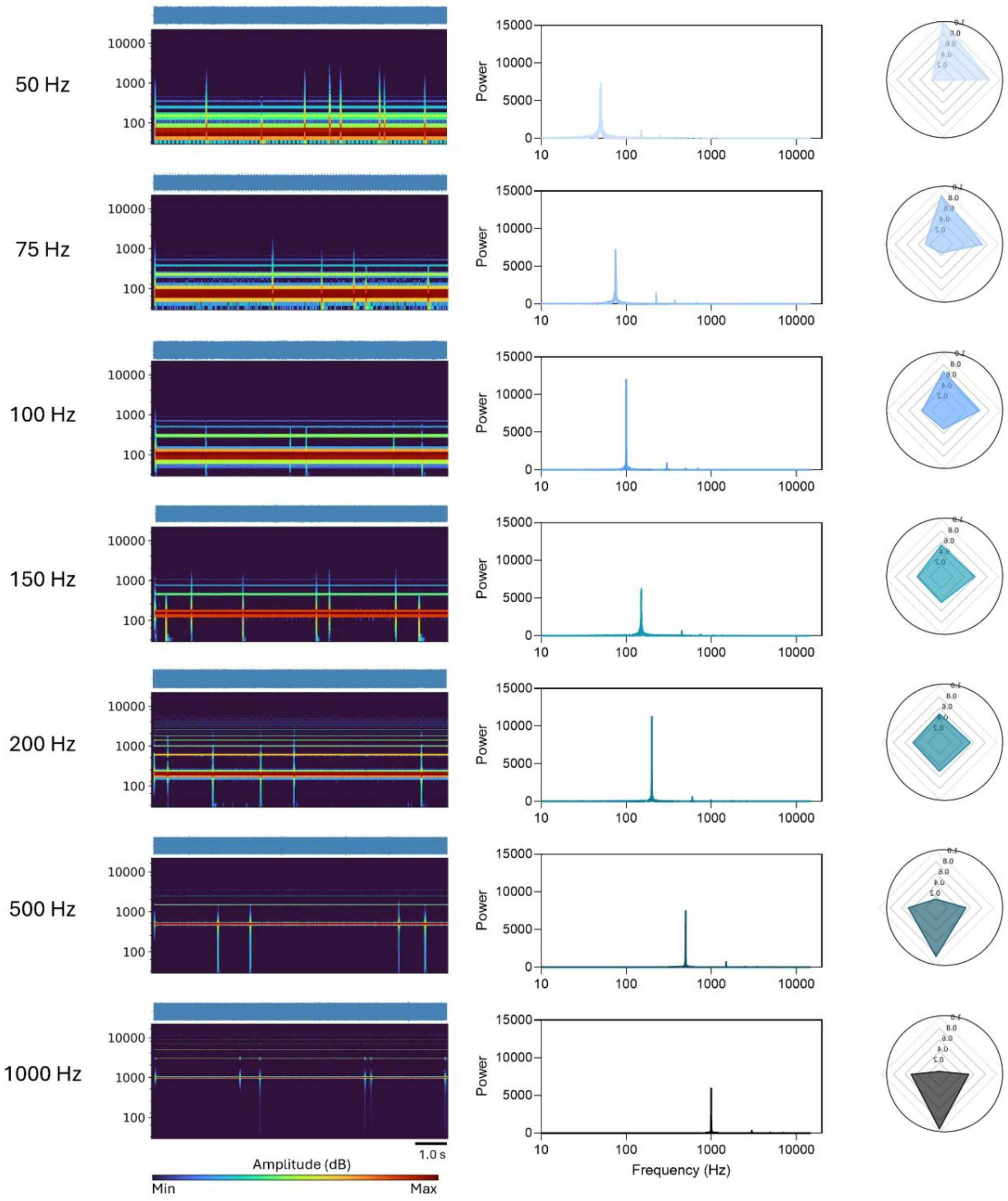


**Figure S9**: Characterization of the basic **Triangle Wave** acoustic input at varying frequencies. A. A heat map of the frequency distribution as a function of time, below the total waveform. Scale bar is 1 s. (Below) a radar plot that is characterized by North, East, South, and West. (North/South) The power at frequencies below 200 Hz (North) and above 200 Hz (South). (East/West) The relative acoustic density below 200 Hz (East), and above 200 Hz (West).

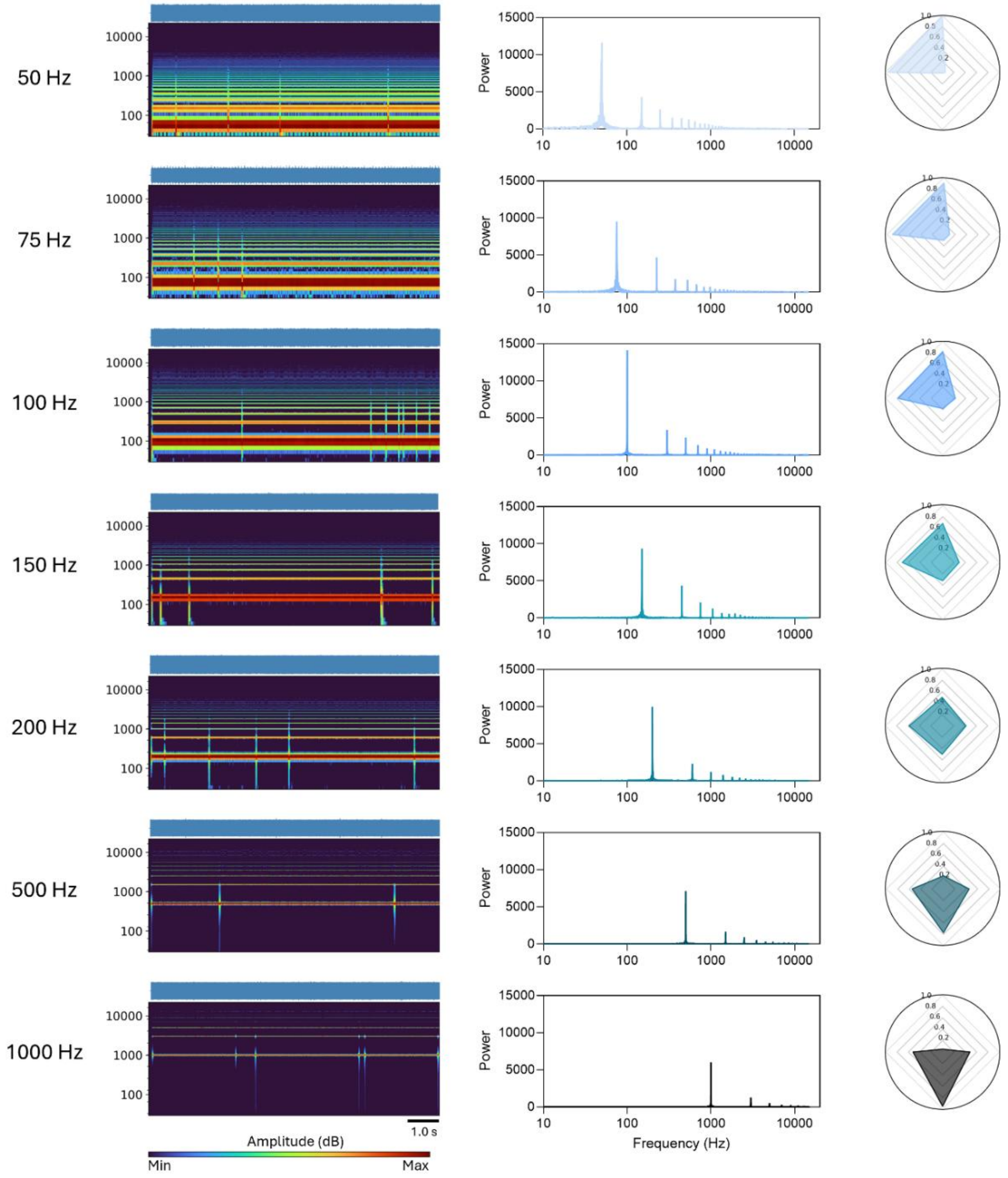


**Figure S10**: Characterization of the basic **Square Wave** acoustic input at varying frequencies. A. A heat map of the frequency distribution as a function of time, below the total waveform. Scale bar is 1 s. (Below) a radar plot that is characterized by North, East, South, and West. (North/South) The power at frequencies below 200 Hz (North) and above 200 Hz (South). (East/West) The relative acoustic density below 200 Hz (East), and above 200 Hz (West).

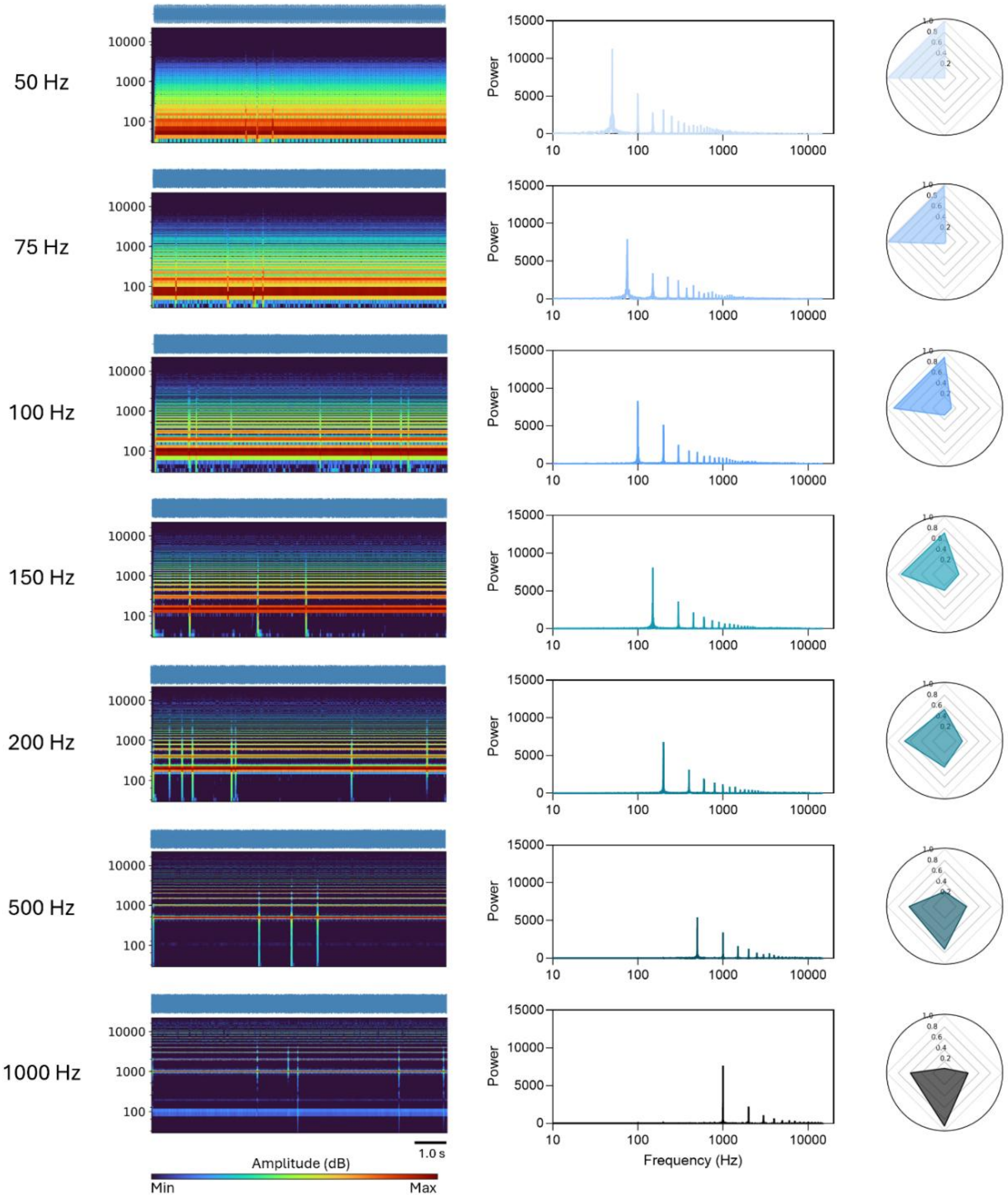


**Figure S11**: Characterization of the basic **Sawtooth Wave** acoustic input at varying frequencies. A. A heat map of the frequency distribution as a function of time, below the total waveform. Scale bar is 1 s. (Below) a radar plot that is characterized by North, East, South, and West. (North/South) The power at frequencies below 200 Hz (North) and above 200 Hz (South). (East/West) The relative acoustic density below 200 Hz (East), and above 200 Hz (West).

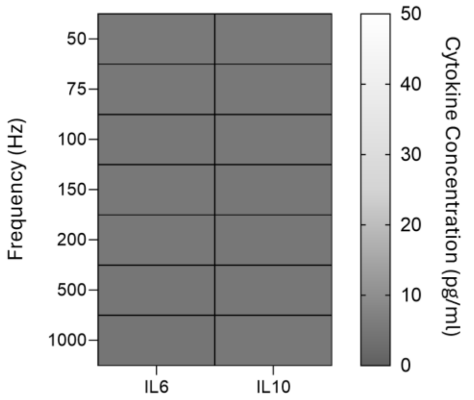


| Condition | M1 | M2 |
|---|---|---|
| 50 Hz | - | - |
| 75 Hz | - | - |
| 100 Hz | - | - |
| 150 Hz | - | - |
| 200 Hz | - | - |
| 500 Hz | - | - |
| 1000 Hz | - | - |

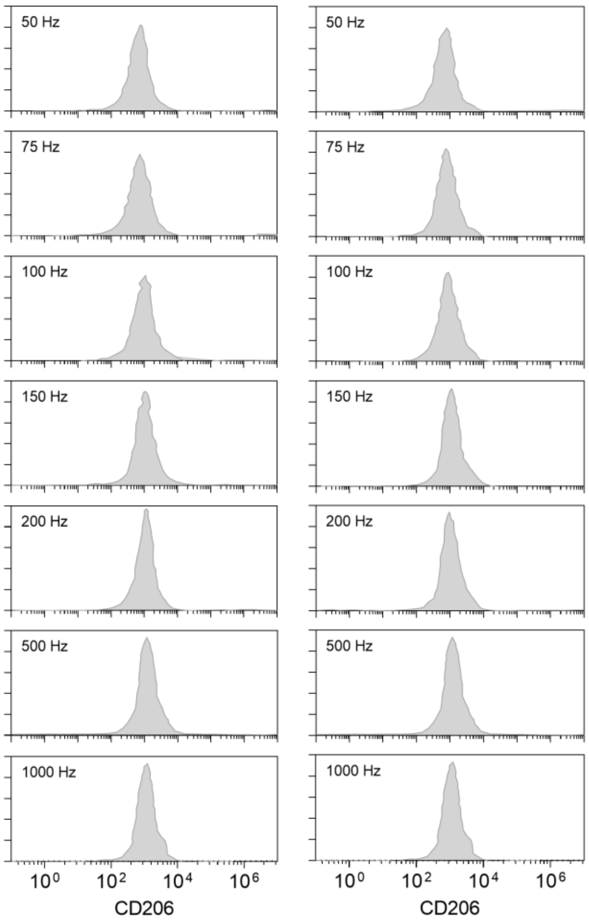


**Figure S12**: Effect of Sine wave frequency on macrophage phenotype through cytokine release studies of IL6, and IL10, and M1 and M2 marker characterization (CD86, and CD206) on the sEVs. The table shows the verification of whether each sample is indicative of M1 or M2 polarized RAW264.7

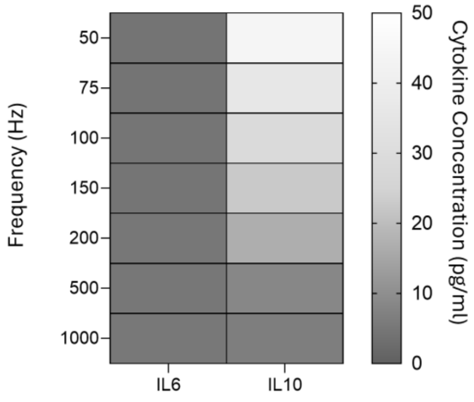


| Condition | M1 | M2 |
|---|---|---|
| 50 Hz | - | ✓ |
| 75 Hz | - | ✓ |
| 100 Hz | - | ✓ |
| 150 Hz | - | ✓ |
| 200 Hz | - | ✓ |
| 500 Hz | - | - |
| 1000 Hz | - | - |

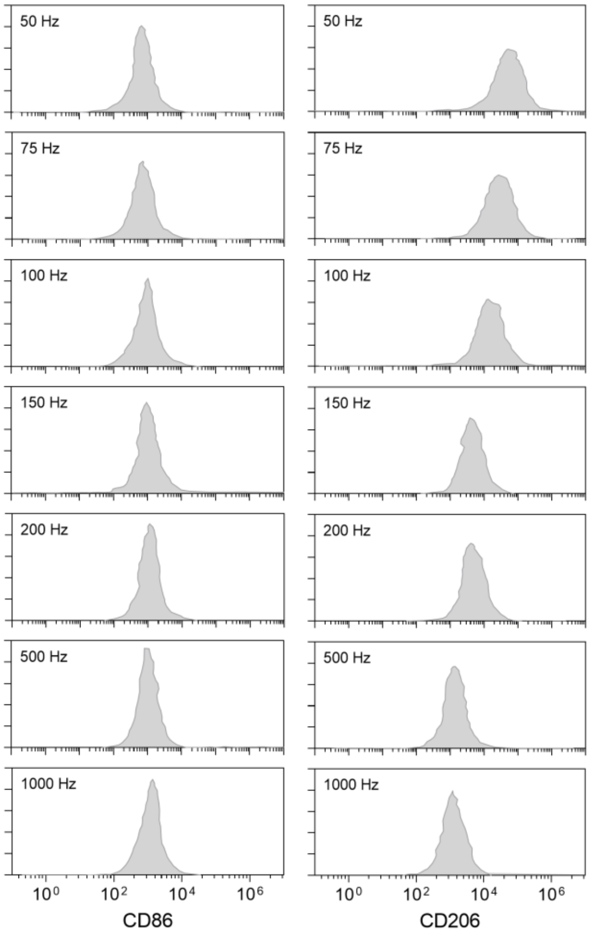


**Figure S13**: Effect of Triangle wave frequency on macrophage phenotype through cytokine release studies of IL6, and IL10, and M1 and M2 marker characterization (CD86, and CD206) on the sEVs. The table shows the verification of whether each sample is indicative of M1 or M2 polarized RAW264.7

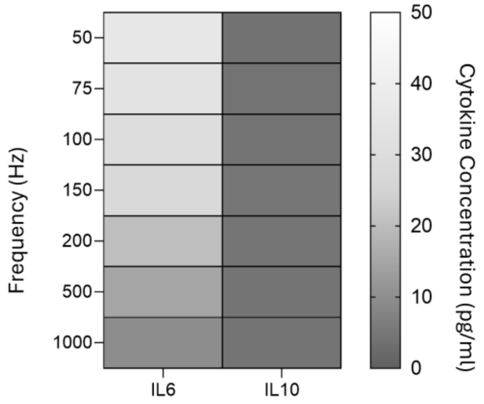


| Condition | M1 | M2 |
|---|---|---|
| 50 Hz | ✓ | - |
| 75 Hz | ✓ | - |
| 100 Hz | ✓ | - |
| 150 Hz | ✓ | - |
| 200 Hz | ✓ | - |
| 500 Hz | - | - |
| 1000 Hz | - | - |

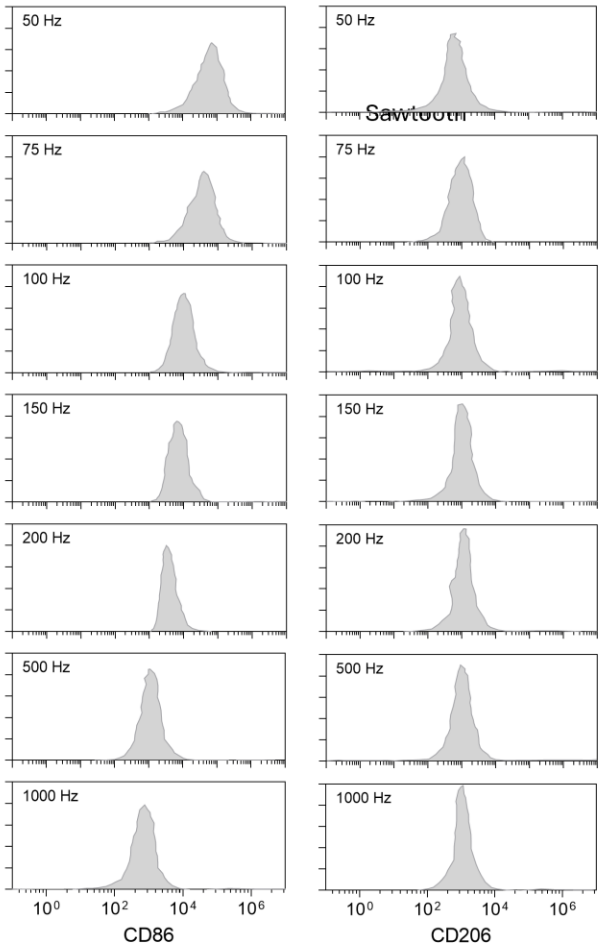


**Figure S14**: Effect of Sawtooth wave frequency on macrophage phenotype through cytokine release studies of IL6, and IL10, and M1 and M2 marker characterization (CD86, and CD206) on the sEVs. The table shows the verification of whether each sample is indicative of M1 or M2 polarized RAW264.7

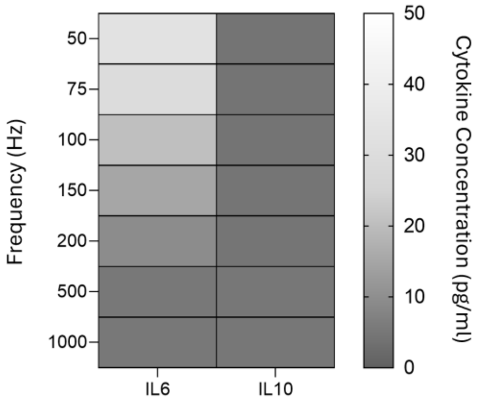


| Condition | M1 | M2 |
|---|---|---|
| 50 Hz | ✓ | - |
| 75 Hz | ✓ | - |
| 100 Hz | ✓ | - |
| 150 Hz | ✓ | - |
| 200 Hz | ✓ | - |
| 500 Hz | - | - |
| 1000 Hz | - | - |

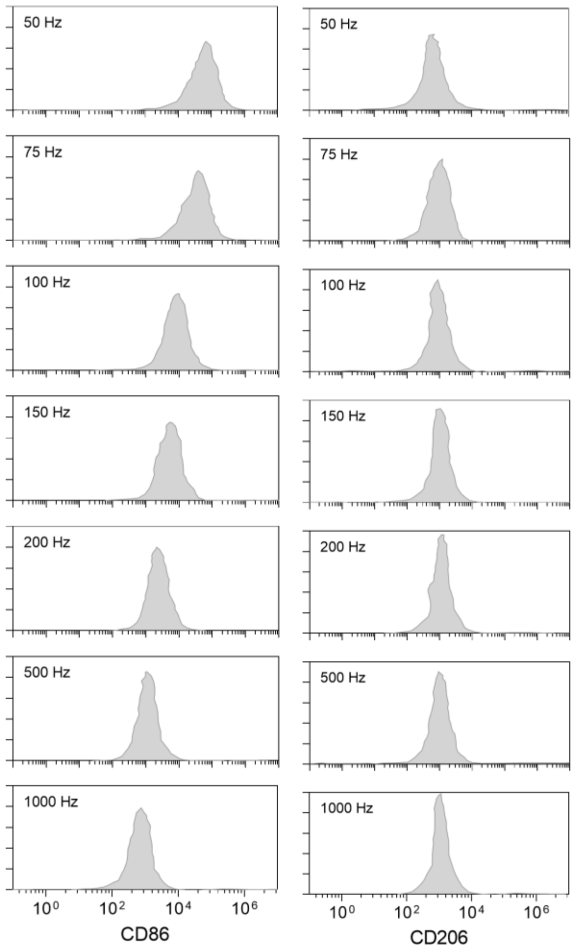


**Figure S15**: Effect of Square wave frequency on macrophage phenotype through cytokine release studies of IL6, and IL10, and M1 and M2 marker characterization (CD86, and CD206) on the sEVs. The table shows the verification of whether each sample is indicative of M1 or M2 polarized RAW264.7

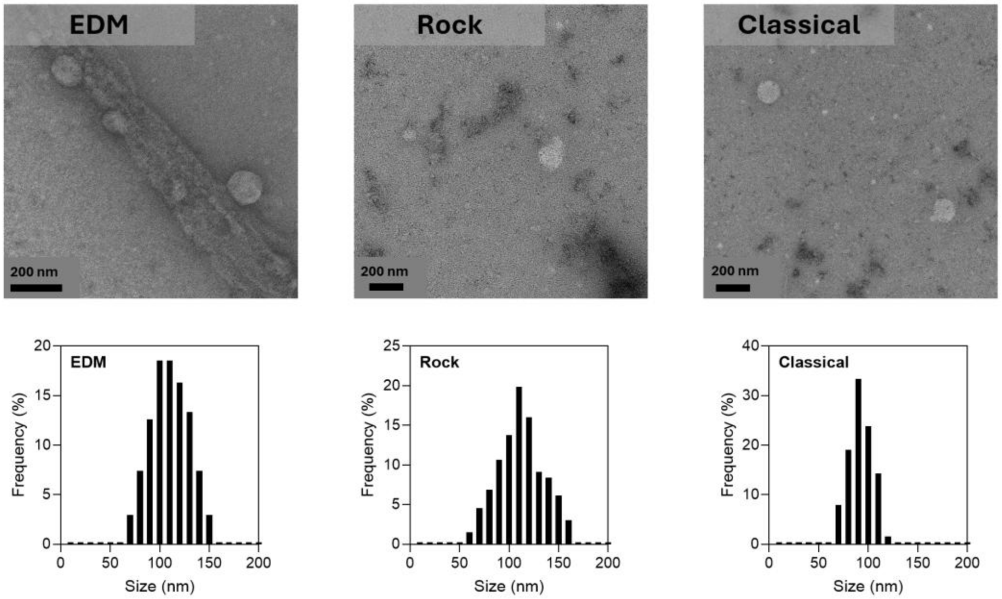


**Figure S16**: TEM imaging of sEVs produced from music genre samples with size distribution analysis (100 particles analyzed).

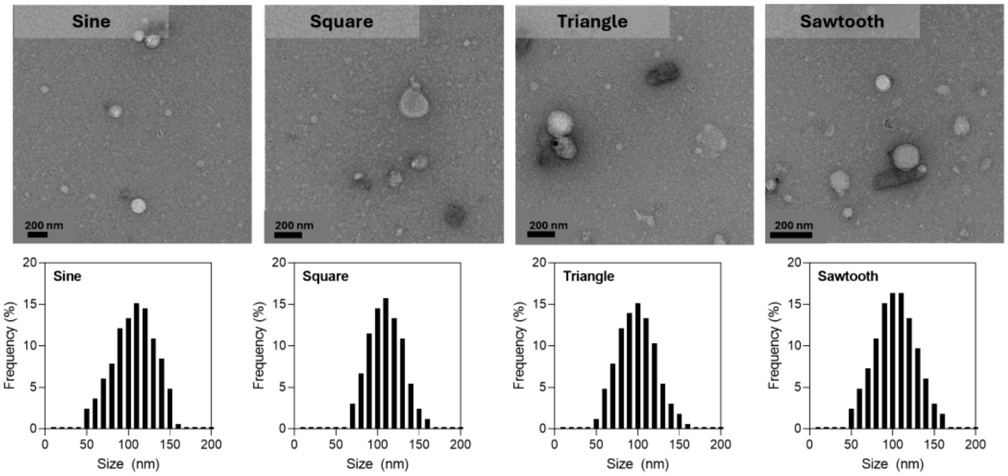


**Figure S17**: TEM imaging of sEVs produced from Various waveform samples (f=100Hz) with size distribution analysis (100 particles analyzed).

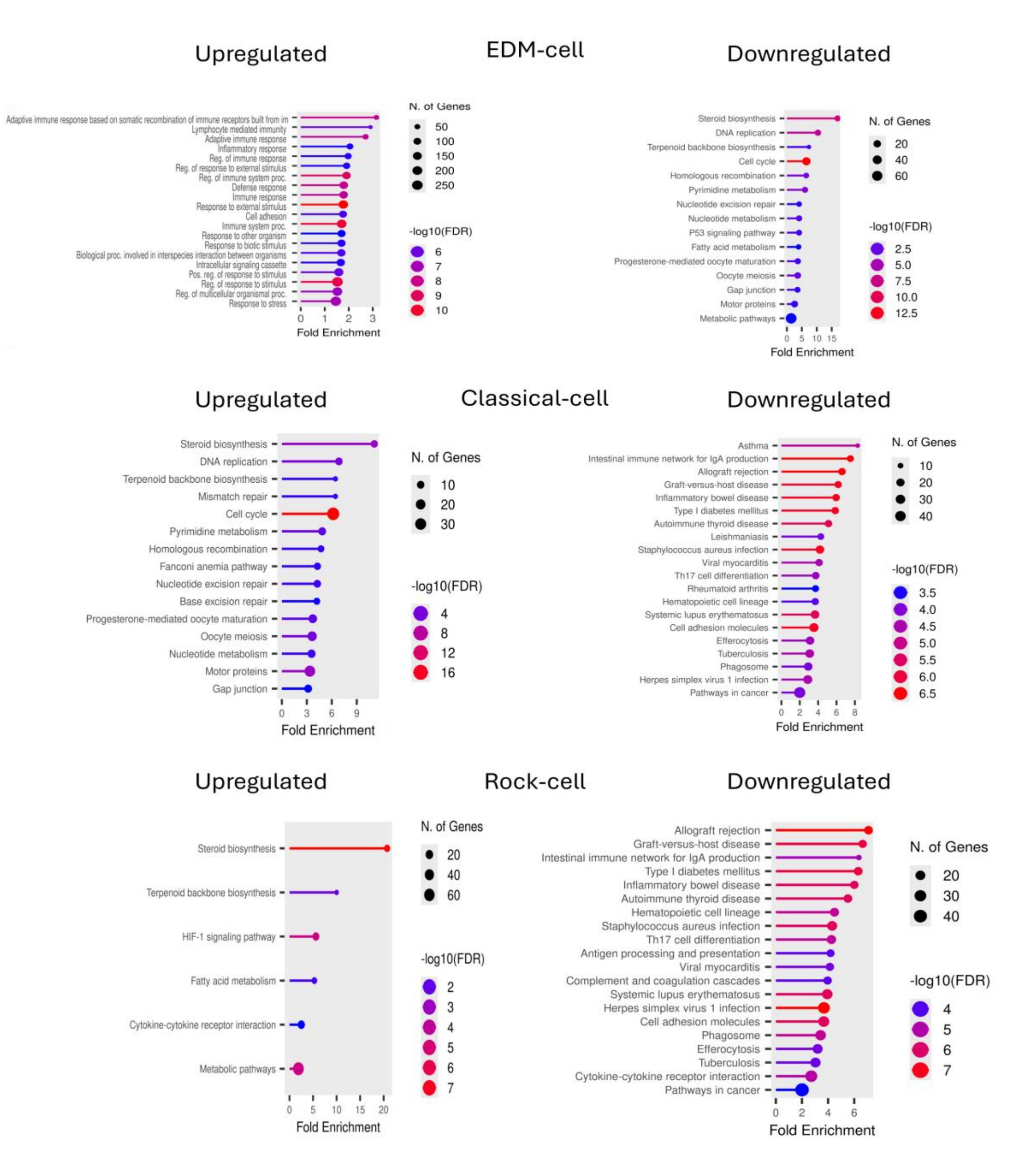


**Figure S18**: KEGG pathway enrichment analysis of RAW264.7 under different musical conditions. (left) Upregulated pathways. (right) Downregulated pathways.

| | sEV markers | | | | | Proinflammatory Cytokines | | | | M1 Macrophage Markers | | | | | | | M0 Macrophage cargo | | M2 Macrophage Markers | | | | | | | MMP | | Collagen subunits | | | |
|---|---|---|---|---|---|---|---|---|---|---|---|---|---|---|---|---|---|---|---|---|---|---|---|---|---|---|---|---|---|---|---|
| Accession # | P41731 | P35762 | P40240 | Q61187 | Q9WU78 | P06804 | P08505 | P10168 | O54824 | P42081 | P18181 | Q18PI6 | O88174 | O89103 | P41272 | P37217 | Q8C8N2 | A2A7Y5 | 035375 | Q6IUU3 | P49615 | Q61830 | QO1149 | P15379 | P29699 | P28862 | P33435 | P08121 | Q5QNQ9 | P39061 | Q64739 |
| 2D CNTR | + | + | + | + | + | - | - | - | - | - | - | - | - | - | - | - | + | + | - | - | - | - | - | - | - | - | - | - | - | - | - |
| PES OFF | + | + | + | + | + | - | - | - | - | - | - | - | - | - | - | - | + | + | - | - | - | - | - | - | - | - | - | - | - | - | - |
| Classical | + | + | + | + | + | - | - | - | - | - | - | - | - | - | - | - | + | + | + | + | + | + | + | + | + | + | + | + | + | + | + |
| Rock | + | + | + | + | + | - | - | - | - | - | - | - | - | - | - | - | + | + | - | - | - | - | - | - | - | - | - | - | - | - | - |
| EDM | + | + | + | + | + | + | + | + | + | + | + | + | + | + | + | + | + | + | - | - | - | - | - | - | - | - | - | - | - | - | - |

**Figure S19**: Proteomics data measured through LC/MS/MS of various sEV samples shows varying hits in cargo associated in M0, M1, and M2 cargo. Moreover, sEVs derived from EDM music have surface markers and cargo protein commonly found in M1 macrophage cells and sEVs. Classical music derived sEVs show the opposite trend with extracellular matrix cargo and surface markers essential for ECM remodeling, making them useful for tissue regeneration applications.

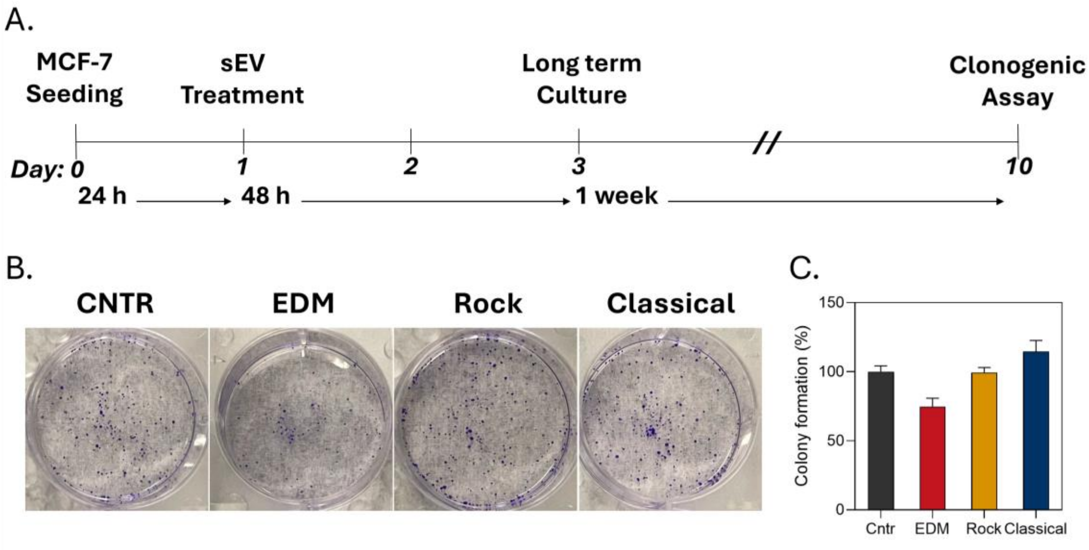


**Figure S20**: Clonogenic assay of MCF7 cells after sEV incubation. A. timeline of clonogenic assay including a 48-hour sEV treatment and a 1 weeklong term culture. B. Picture results of the clonogenic assay from stimuli derived sEVs. C. quantitative analysis of colony formation.

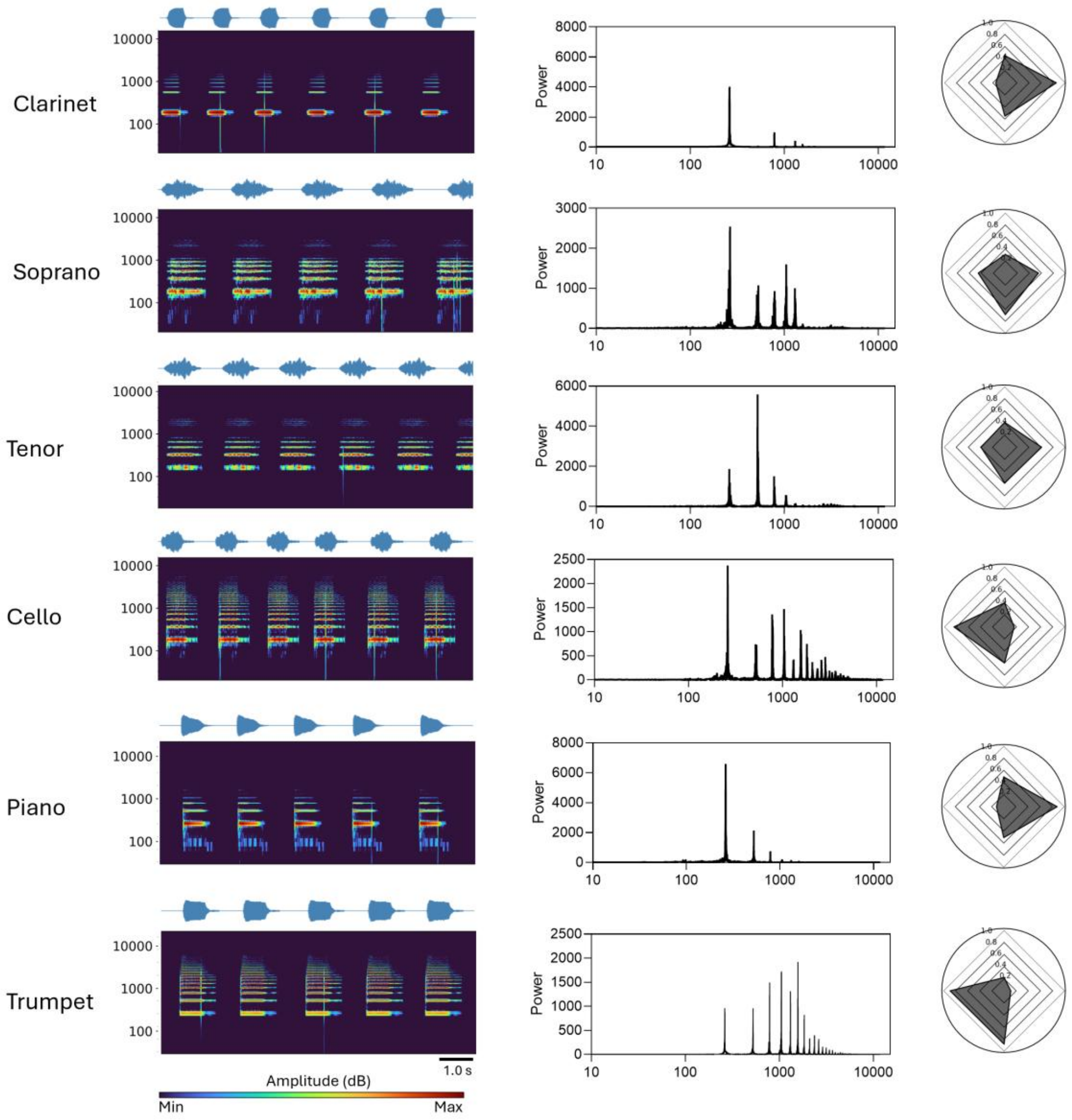


**Figure S21**: Characterization of the **Instrument acoustic** input at varying frequencies. A. A heat map of the frequency distribution as a function of time, below the total waveform. Scale bar is 1 s. (Below) a radar plot that is characterized by North, East, South, and West. (North/South) The power at frequencies below 200 Hz (North) and above 200 Hz (South). (East/West) The relative acoustic density below 200 Hz (East), and above 200 Hz (West).

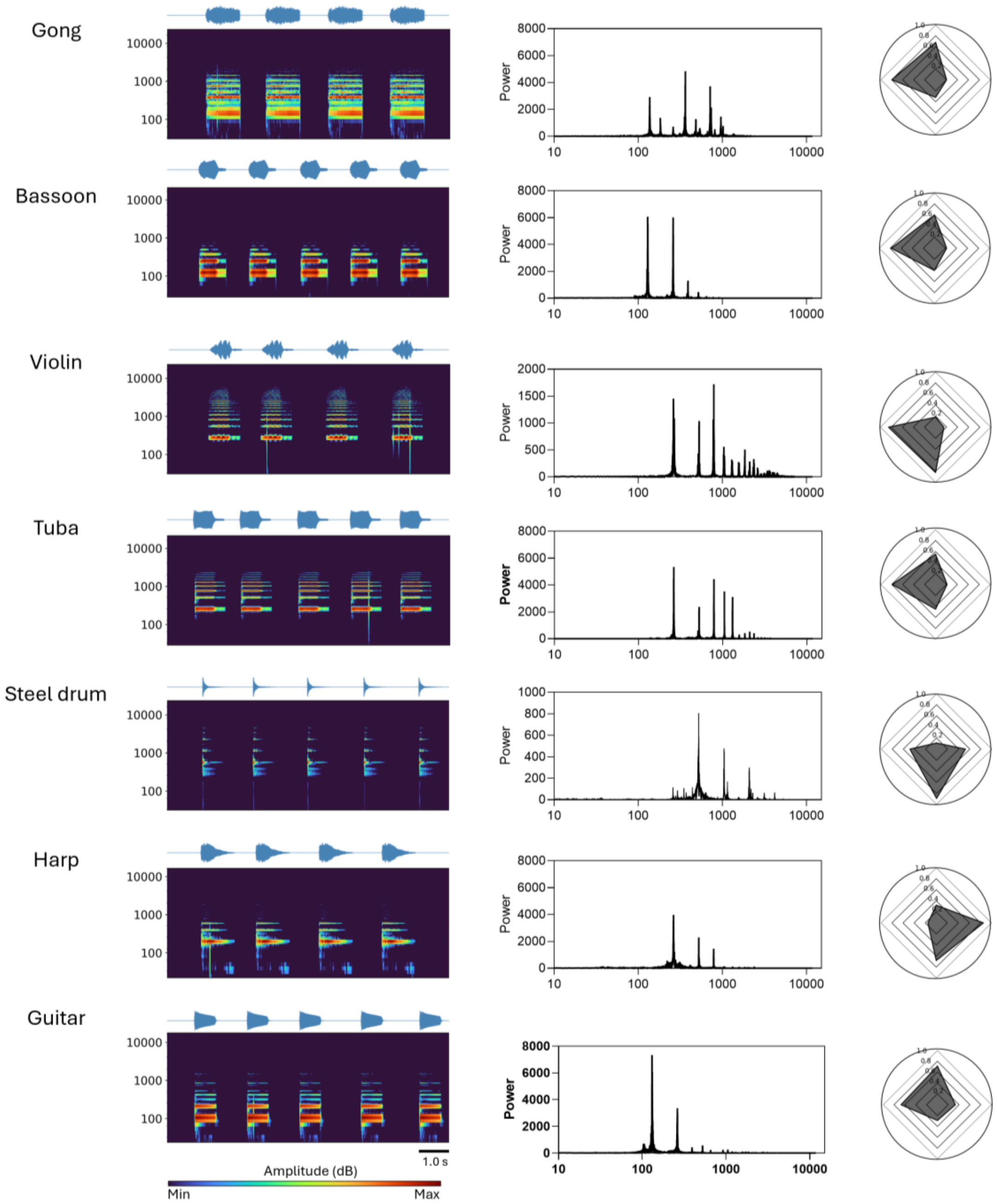


**Figure S22**: Characterization of the basic **instrument acoustic** input at varying frequencies. A. A heat map of the frequency distribution as a function of time, below the total waveform. Scale bar is 1 s. (Below) a radar plot that is characterized by North, East, South, and West. (North/South) The power at frequencies below 200 Hz (North) and above 200 Hz (South). (East/West) The relative acoustic density below 200 Hz (East), and above 200 Hz (West).

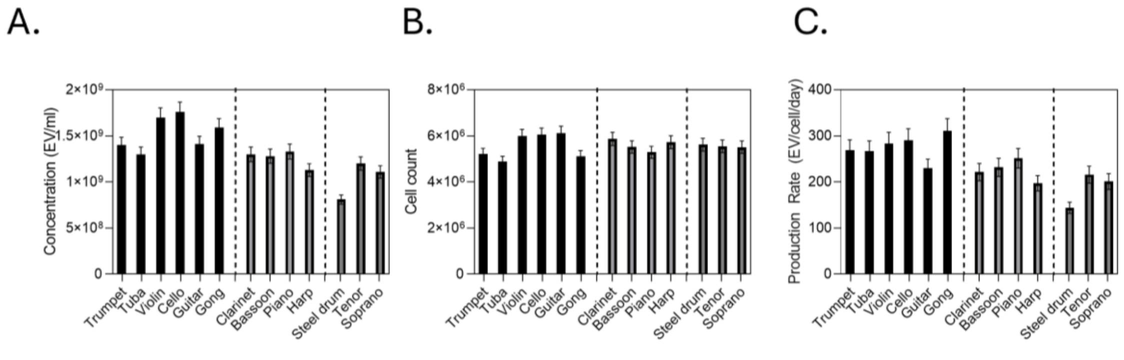


**Figure S23**: sEV production testing and verification of the effect on Instrument acoustic input on sEV production at various frequency inputs. Each instrument was played at a C4 note (255 Hz fundamental frequency) A. Concentration measurements using NTA (N=5). B. Cell count measurements using manual cell counting after detaching the cells from the PES material (N=5). C. Production rate calculation from the previous measurements at 24-hour incubation.

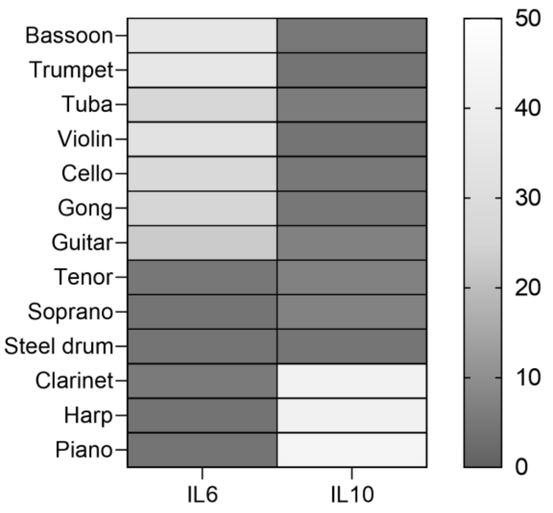


| Instrument | M1 | M2 |
|---|---|---|
| **Bassoon** | ✓ | - |
| **Trumpet** | ✓ | - |
| **Tuba** | ✓ | - |
| **Violin** | ✓ | - |
| **Cello** | ✓ | - |
| **Gong** | ✓ | - |
| **Guitar** | - | - |
| **Tenor** | - | - |
| **Soprano** | - | - |
| **Steel Drum** | - | - |
| **Clarinet** | - | ✓ |
| **Harp** | - | ✓ |
| **Piano** | - | ✓ |

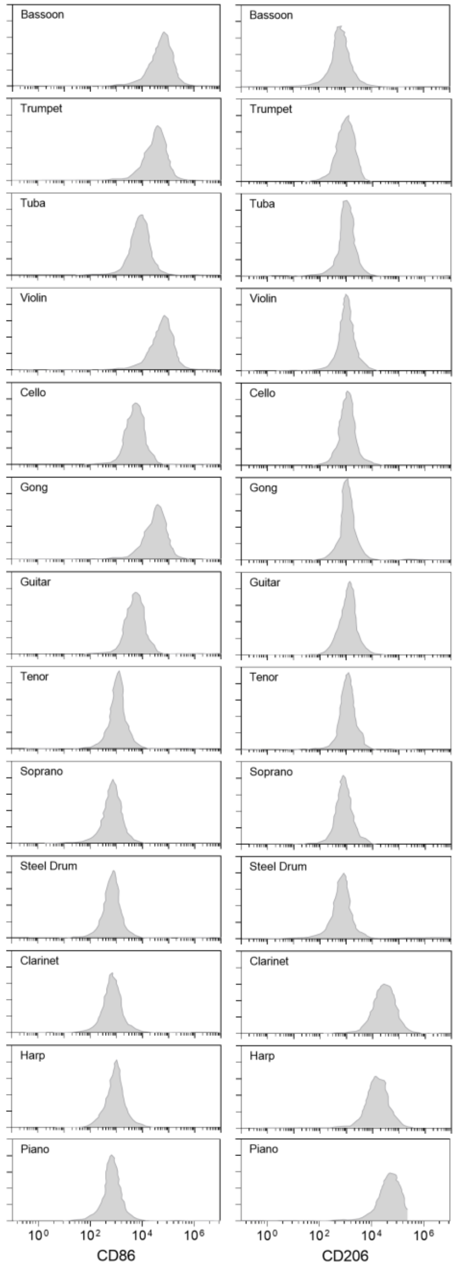


**Figure S24**: Effect of instrument input at 255 Hz fundamental frequency on macrophage phenotype through cytokine release studies of IL6, and IL10, and M1 and M2 marker characterization (CD86, and CD206) on the sEVs. The table shows the verification of whether each sample is indicative of M1 or M2 polarized RAW264.7.

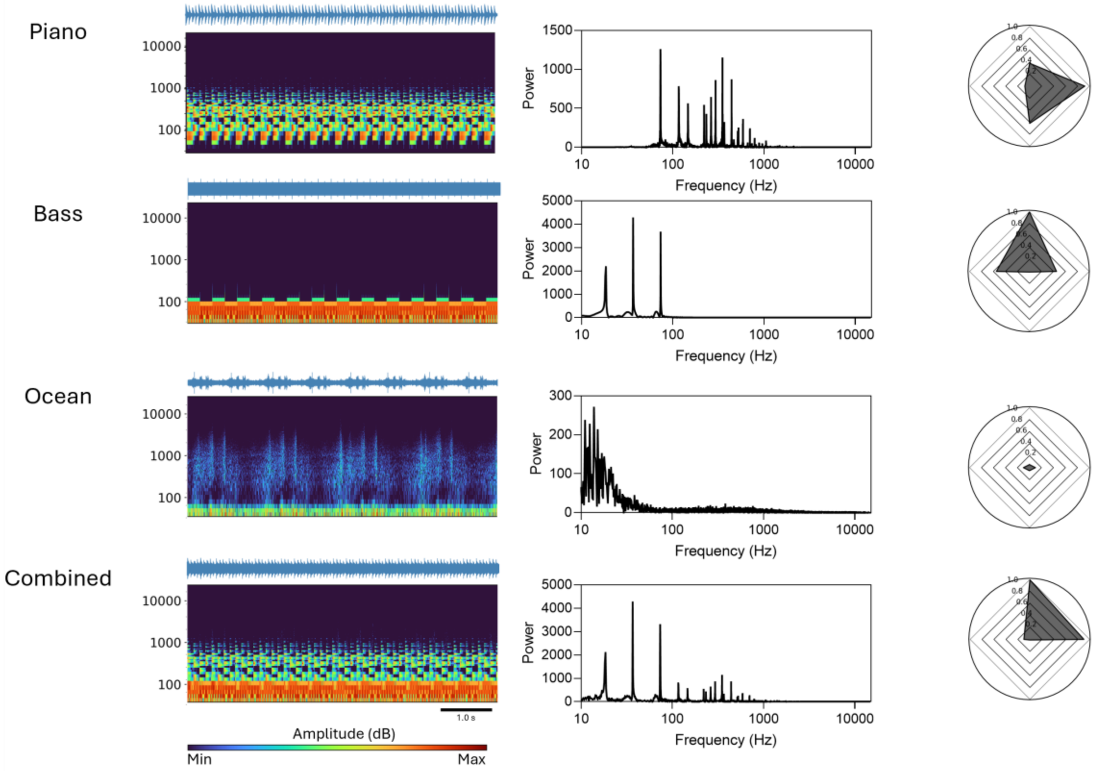


**Figure S25**: Characterization of each component of the regeneration music input. A. A heat map of the frequency distribution as a function of time, below the total waveform. Scale bar is 1 s. (Below) a radar plot that is characterized by North, East, South, and West. (North/South) The power at frequencies below 200 Hz (North) and above 200 Hz (South). (East/West) The relative acoustic density below 200 Hz (East), and above 200 Hz (West).

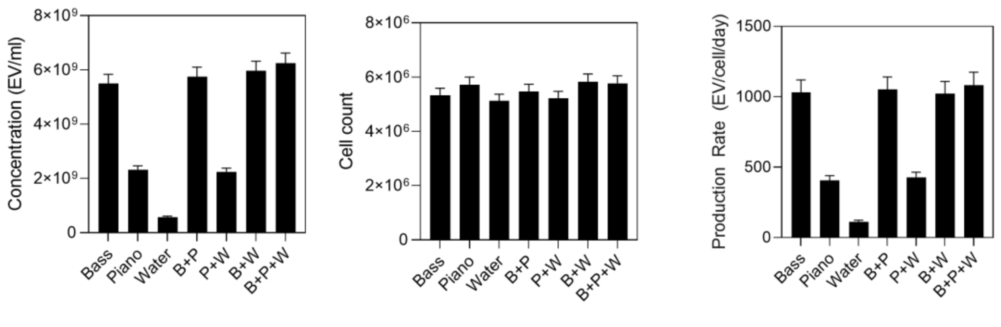


**Figure S26**: sEV production testing and verification of regeneration component on sEV production. A. Concentration measurements using NTA (N=5). B. Cell count measurements using manual cell counting after detaching the cells from the PES material (N=5). C. Production rate calculation from the previous measurements at 24-hour incubation.

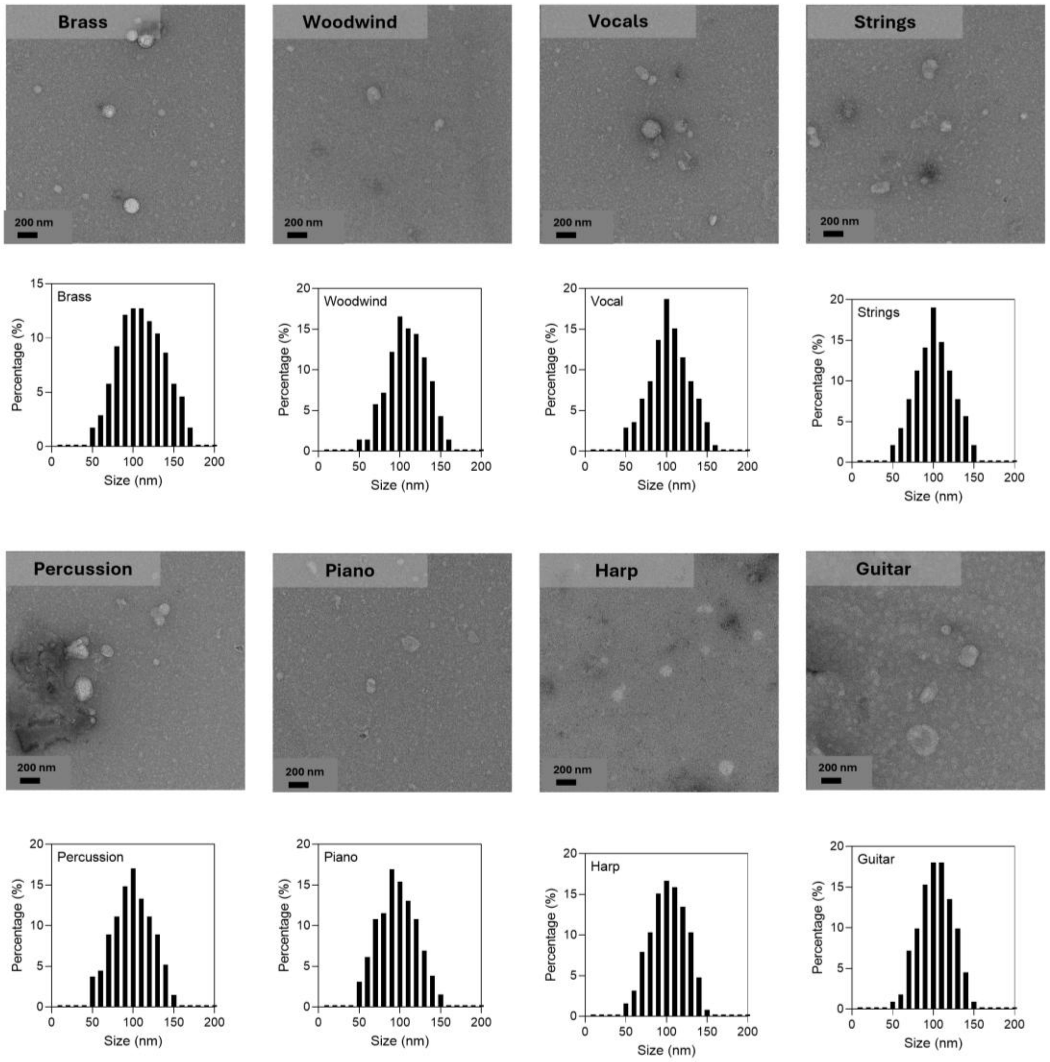


**Figure S27**: TEM imaging of sEVs produced from categorized instrument samples with size distribution analysis (100 particles analyzed).

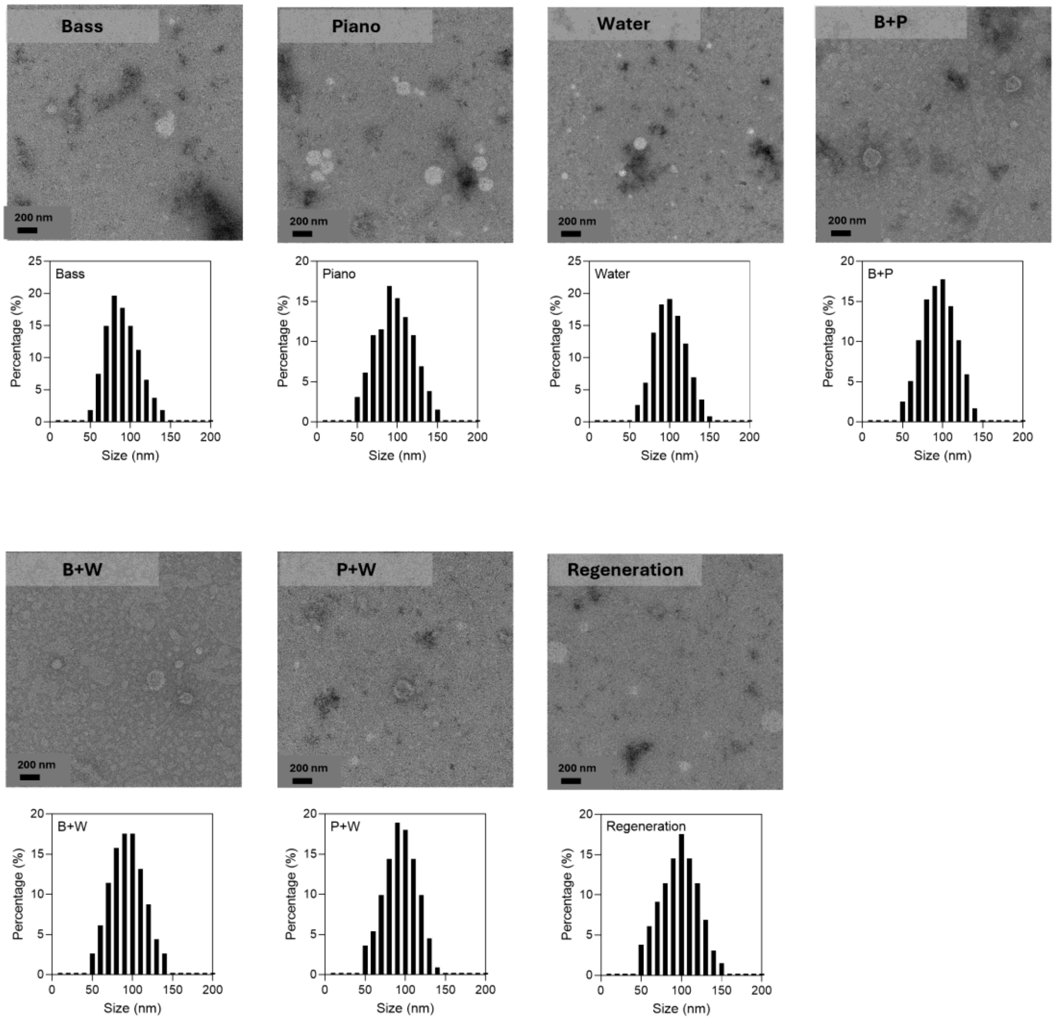


**Figure 28**: TEM imaging of sEVs produced from engineered regeneration components with size distribution analysis (100 particles analyzed).
